# Early peripheral immune signaling precedes tau elevation and blood–brain barrier disruption in Alzheimer’s disease

**DOI:** 10.64898/2026.04.02.716122

**Authors:** Aaron Burberry, Penelope Benchek, Mark Lowe, Wanyong Shin, Blake McCourt, Quintasha Beamon, Shinjon Chakrabarti, Sara Ramaiah, Elizabeth Woidke, Maria Khrestian, Holden Maecker, Lynn M. Bekris, Stephen Rao, Daniel Ontaneda, James B. Leverenz, William Bush, Jagan A. Pillai

## Abstract

**Abstract:** Neuroinflammation, along with amyloid beta (Aβ) deposition, phospho-tau (ptau) accumulation, blood–brain barrier (BBB) disruption, and cognitive decline are recognized components of Alzheimer’s disease (AD). However, the timing and nature of peripheral immune changes across AD biological and clinical stages remain poorly understood. Here we performed mass cytometry profiling of whole blood and cerebrospinal fluid (CSF) immune cells from 351 human samples across two independent clinical cohorts spanning the AD continuum. We identify coordinated peripheral immune signaling signatures that emerge during preclinical stage of AD and precede significant elevation of plasma ptau217, CSF ptau181 and BBB disruption measured by dynamic contrast–enhanced magnetic resonance imaging (DCE-MRI). AD-enriched immune features, including increased phospho-Akt signaling in naï ve T killer cells and phospho-PLCγ2 signaling in granulocytes, were not observed in patients with Frontotemporal lobar degeneration or treatment-naï ve multiple sclerosis. Furthermore, these immune signaling states could be induced in healthy donor immune cells following exposure to plasma or CSF from individuals with AD, indicating that circulating factors can drive these peripheral immune alterations. Together, our findings demonstrate that dynamic peripheral immune state changes arise early in AD and precede canonical biomarker and vascular changes, highlighting immune signaling pathways as potential targets for early therapeutic intervention.

**Graphical Abstract:** Early peripheral immune signaling precedes tau elevation and blood-brain barrier disruption in Alzheimer’s disease.

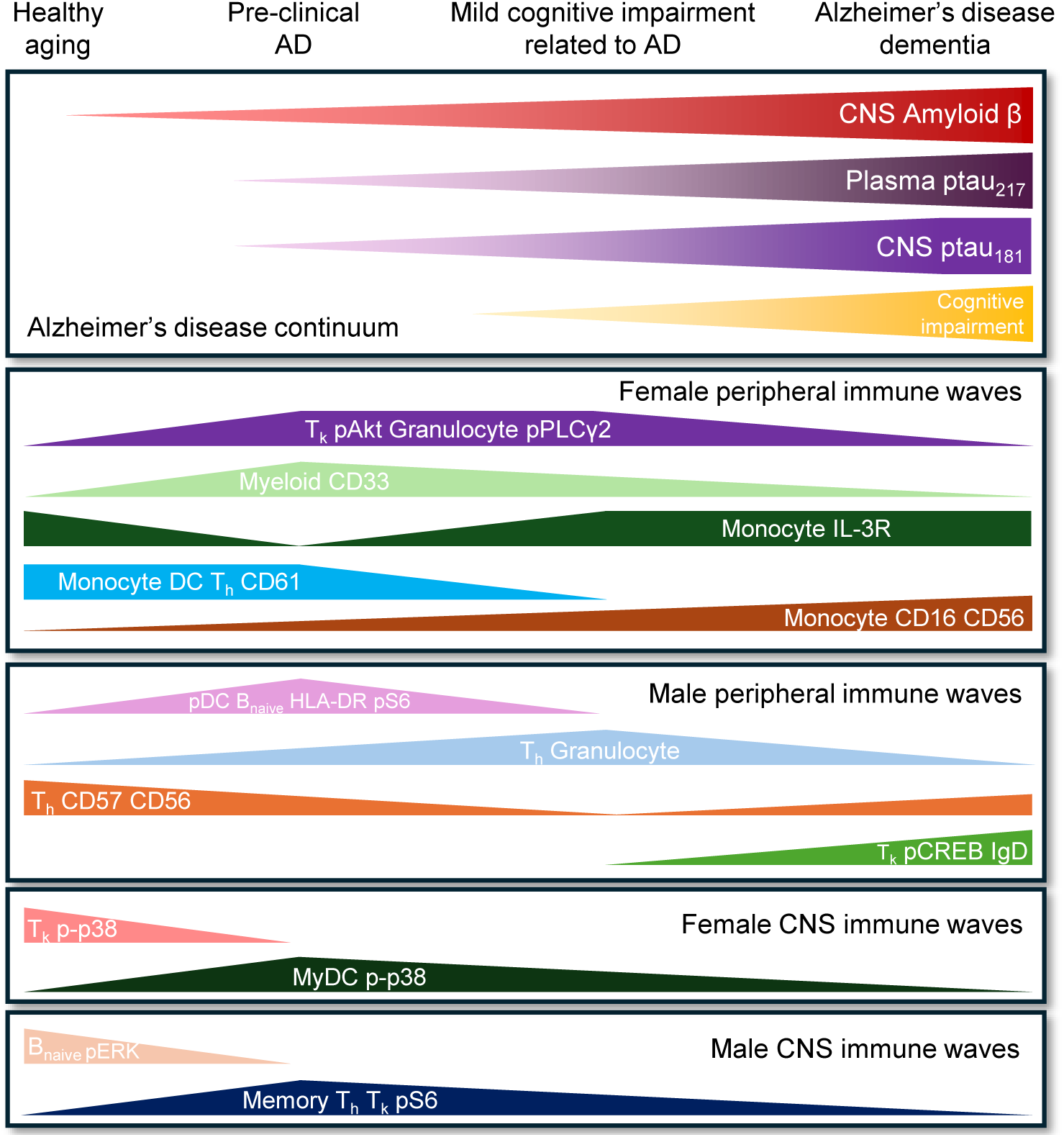

## Introduction

Alzheimer’s disease, the most common etiology of dementia worldwide, is classically defined by neuropathological hallmarks of Aβ plaques and neurofibrillary tau tangles. However a wealth of evidence from animal models, human neuropathologic evaluation, genome-wide association studies, and clinical studies highlights immune-related dysregulation as a key contributor to AD pathophysiology^1,2^. The contribution of brain-resident microglia at later stages of AD is well-characterized^3–5^, yet the timeline of peripheral immune cell changes and their effector molecules in AD is poorly understood, with some evidence suggesting that peripheral inflammatory proteins are elevated after β amyloid and ptau pathologic markers^6^. Furthermore, the relationship between the peripheral immune alterations and changes in CSF immune populations, blood brain barrier (BBB) integrity and cognitive impairment remains unclear.

Monocytes participate in the clearance of Aβ aggregates from peripheral vasculature at the mild cognitive impairment (MCI) stage of AD^7,8^. Yet there is uncertainty as to whether the abundance and function of these circulating innate immune cells become altered before the onset of cognitive impairment, in the preclinical stage of AD^7–10^. In the adaptive immune compartment, T cells with an effector memory phenotype that express high levels of CD45RA (T_EMRA_) are reported to be enriched in peripheral blood and CSF of individuals across pre-clinical-AD^11^, MCI stage^11–13^ and during late-stage AD dementia^12,13^, whereas other studies have found no evidence for T_EMRA_ enrichment in peripheral blood of AD patients despite profiling the requisite surface markers^14^. Beyond changes in specific immune cell numbers, stimulation of peripheral^14^ or CNS-derived^15^ immune cells with inflammatory moieties have uncovered AD-related gene expression programs, yet efforts to identify resting immune cell signatures in the periphery or CNS that define stages of AD have yet to be reported. To address these gaps, and to test the hypothesis that alterations in immune cell states would accompany AD biological and clinical stages, we profiled innate and adaptive immune networks simultaneously in peripheral blood and CSF across distinct biological and clinical stages of AD^16^.

BBB integrity is well appreciated to promote healthy aging and is noted with AD progression^17^. Protein signatures in the plasma and CSF correlate with the extent of BBB breakdown as well as cognitive impairment related to AD and other neurodegenerative disorders^18–21^. Yet debate continues on whether peripheral immune dysregulation in AD requires BBB breakdown and if the diffusion of factors or cells into the CNS precedes cognitive impairment related to AD^22–25^. Knowledge of which immune cell types herald worsening disease severity has important implications for evaluating rate of disease progression and potential stage-specific therapeutic interventions in AD. This effort could also provide insight into therapeutic opportunities and molecular mechanisms underlying cognitive resilience in the healthy aging population. To address these questions, we applied mass cytometry time of flight (CyTOF) to unstimulated peripheral blood and CSF cells derived from individuals that underwent concordant dynamic contrast enhanced magnetic resonance imaging (DCE-MRI) to comprehensively characterize peripheral and CNS immune cell states and their relationship with BBB permeability across stages of AD. Key findings related to peripheral immune activation were replicated by re-analysis of an independent PBMC CyTOF dataset^13^. Together, this allowed us to define the temporal order and compartment-specific nature of immune activation in an AD stage-dependent manner.

## Results

To understand the relationship between immune activation states in the periphery and CNS across clinical stages of AD, we used mass cytometry to analyze immune cells from two cohorts. In a discovery cohort at the Cleveland Clinic Center for Brain Health, Cleveland, OH, we analyzed paired whole blood and CSF cells from age-matched individuals (70.8±3.7 years) including cognitively normal controls (CN, n=27 blood samples; n=24 CSF samples), cognitively normal individuals at the pre-clinical stage of AD (CN Aβ+, defined by a reduction of the Aβ_42/40_ ratio in CSF, n=16 blood samples; n=15 CSF samples), mild cognitive impairment stage of AD (MCI, n=33 blood samples, n=30 CSF samples) and dementia stage of AD (AD, n=23 blood samples, n=21 CSF samples) (**Fig 1a** and **Table 1**). To validate key results, we reanalyzed a mass cytometry dataset^13^ of PBMC derived from age-matched subjects (62.7±7.3 years) from the Amsterdam Dementia Cohort^26^. PBMC donors had baseline diagnoses of subjective cognitive decline (SCD, n=33), MCI (n=42) or AD (n=59) (**Table 1**). We reasoned that consistent findings between the two cohorts may represent disease-related peripheral immune changes that are robust to methodologic and demographic covariates, whereas the differences that we observe may reflect variation in immune cell processing methods for CyTOF, the age range of donors, the proportion of *APOE ε4* risk alleles we sampled, the degree of cognitive impairment represented by clinical cases, the frequency of comorbidities such as cardiovascular disease and diabetes and baseline levels of systemic inflammation^27^. Importantly, to control for immune alterations that were unrelated to AD pathophysiology, participants were screened to exclude active anti-inflammatory medication use other than aspirin or a history of bacterial infection, viral infection, autoimmune disease, malignancy or hospitalization at least three months prior to evaluation.

**Fig. 1.**
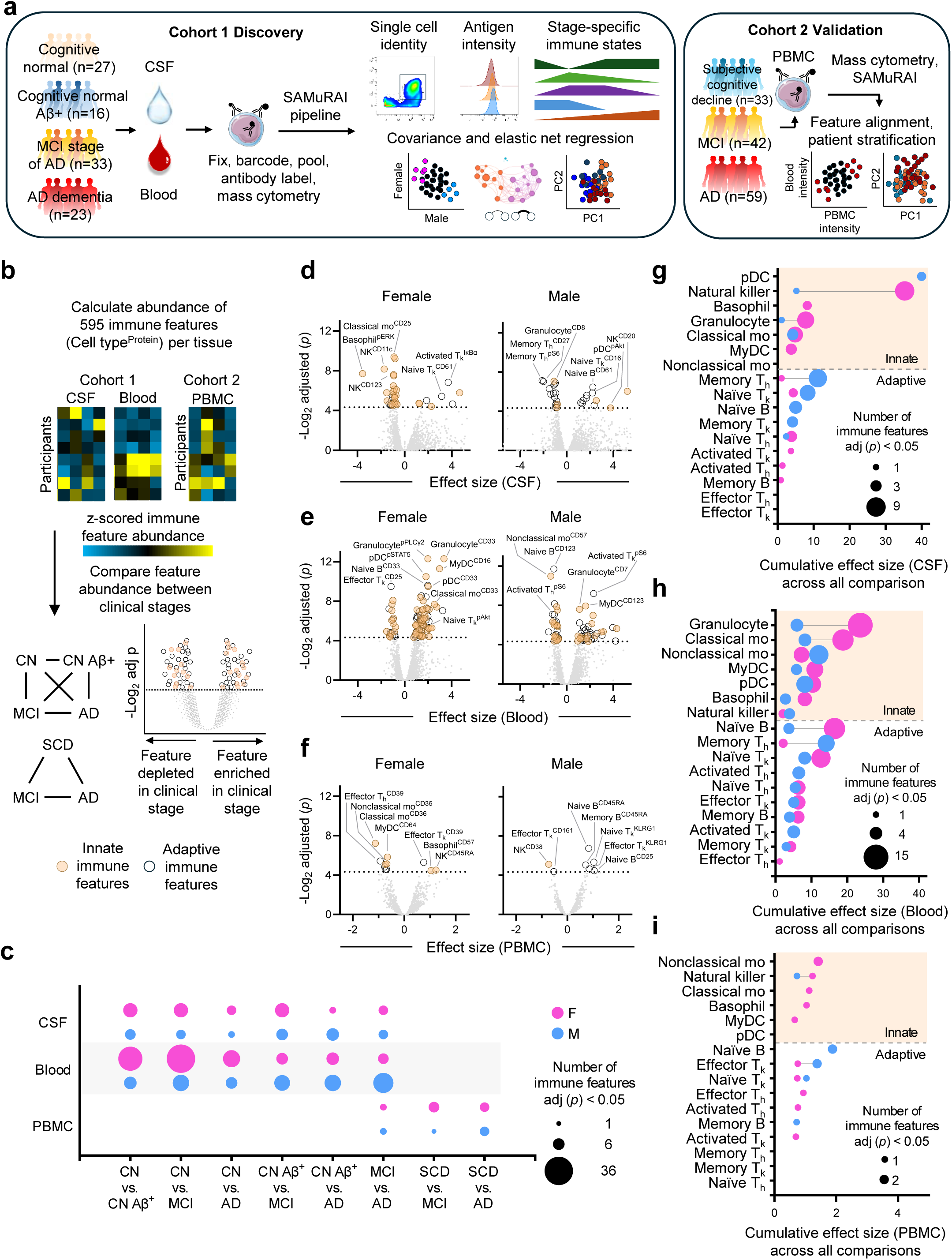
Mass cytometry reveals immune changes in cerebral spinal fluid and blood across clinical stages of Alzheimer’s disease. **a**, Mass cytometry profiling summary covering 99 whole blood samples, 90 cerebral spinal fluid (CSF) samples and 134 PBMC samples from clinically diagnosed participants that visited the Cleveland Clinic (Cohort 1 Discovery) or the Amsterdam Demetia Cohort 2 validation. **b,** Strategy to derive z-score normalized immune features and compare feature abundance between AD stages. Innate immune cell (brown circles) and adaptive immune cell features (open black circles). **c**, The number of differentially abundant immune features in females (pink) and males (blue) were plotted for each comparison. Two-way ANOVA with Tukey multiple comparison. adjusted p-value < 0.05. **d-f**, Volcano plots of differentially abundant immune features in CSF (**d**), blood (**e**) or PBMC (f) datasets. **g-i,** Bubble plots show the number of differentially abundant immune features and cumulative effect size across all comparisons described in (**b)** for each cell type within CSF (**g**), blood (**h**) and PBMC (**i**).

**Table 1.**
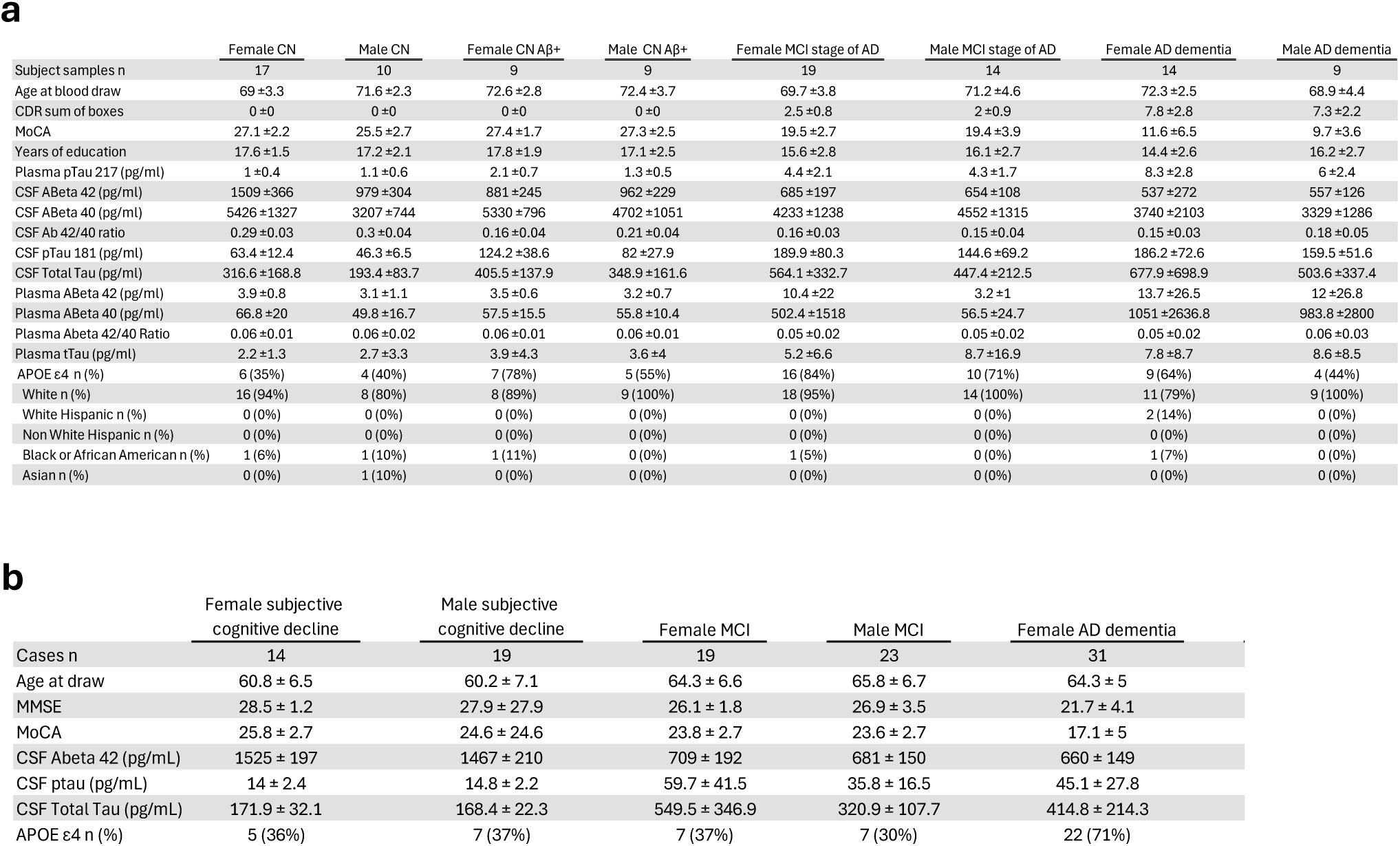
Clinical characteristics of the study cohorts. **a,** Cohort 1, Cleveland Clinic. **b,** Cohort 2, Amsterdam Dementia Cohort^13,26^. Values are reported as mean ± standard deviation.

To enable direct comparison of immune cell abundance and activation state in the periphery and CNS, we developed SAMuRAI (Single-cell Analysis of Multiparametric Regularized Antigen Intensity), a pipeline that distills CyTOF data into normalized proteomic features per sample. The computational approach begins with the identification of major innate and adaptive immune cell populations in human biofluids^28^ at single cell resolution (**Extended data Fig 1a-c** and **Table 2**). By excluding rare populations that can be visualized with other tools^29–31^, this approach reproducibly identified 17 immune populations in blood, CSF and PBMC fractions with the exception of granulocytes which are excluded during the isolation of PBMC (**Extended data Fig 1a-c** and **Table 3** and **Table 4**). To interrogate cellular states, we derived the geometric mean intensity of each antigen in the panel for each gated population. This resulted in 595 immune features (35 antigens x 17 populations) per blood or CSF sample and 592 immune features (37 antigens x 16 populations) per PBMC sample (**Extended data Fig 1e-g**). Among these, 304 immune features (19 antigens x 16 populations) were conserved between the blood and PBMC datasets. To control for run-to-run variability and to enable harmonization between the Cleveland and Amsterdam datasets, each immune feature was batch normalized into a z-score. By comparing the trajectory of immune features across individuals scored for cognition, memory and DCE-MRI measures of BBB permeability, we identified concordant immune features and modules that were engaged in an AD-stage specific manner and uncovered key nodes of influence within the immune network.

**Table 2.**
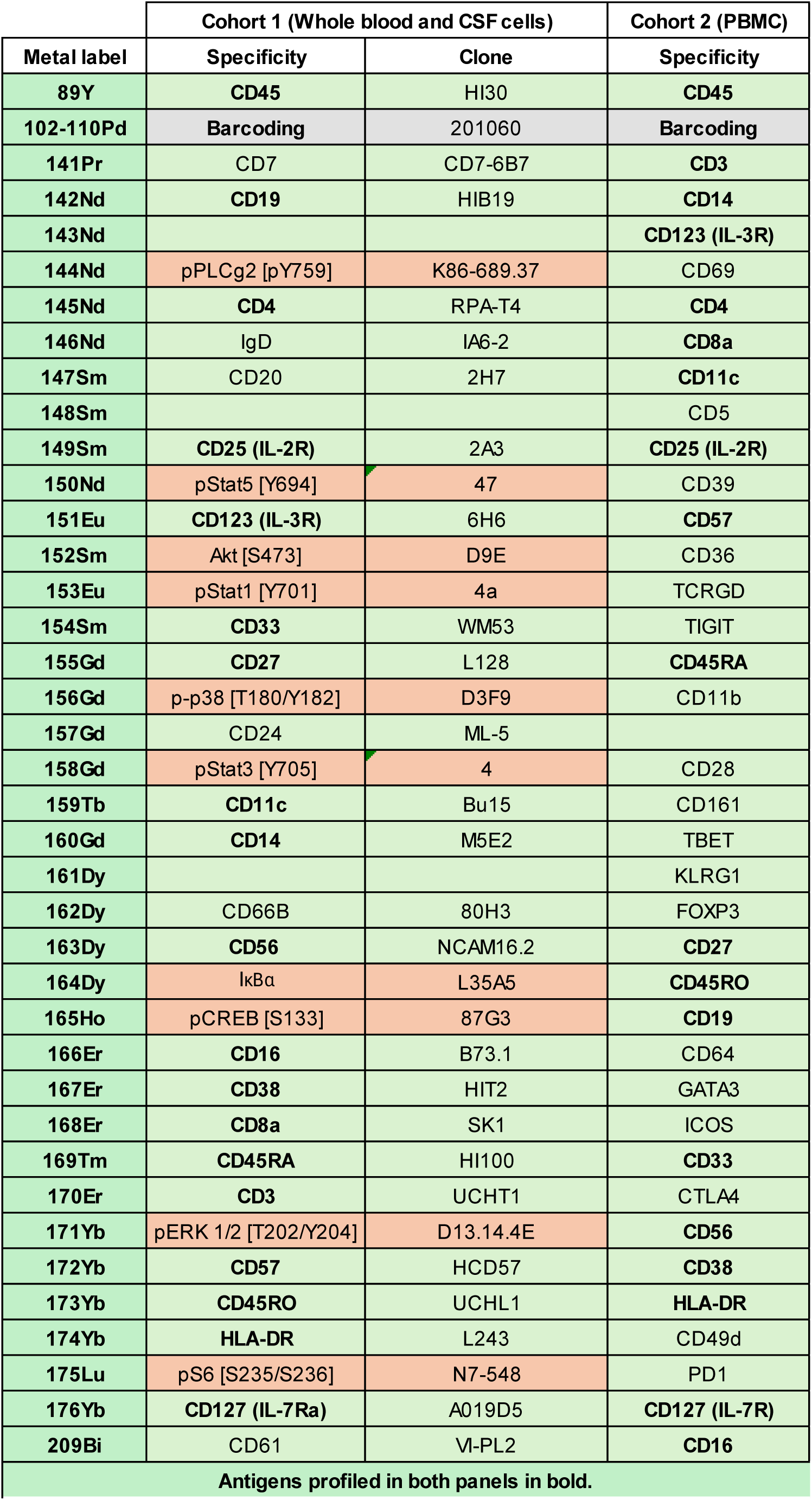
Mass cytometry antibody panels. Table of mass tag for each antibody, the antigens targeted, and antibody clones. Custom conjugates performed by Stanford Human Immune Monitoring Center.

**Table 3.**
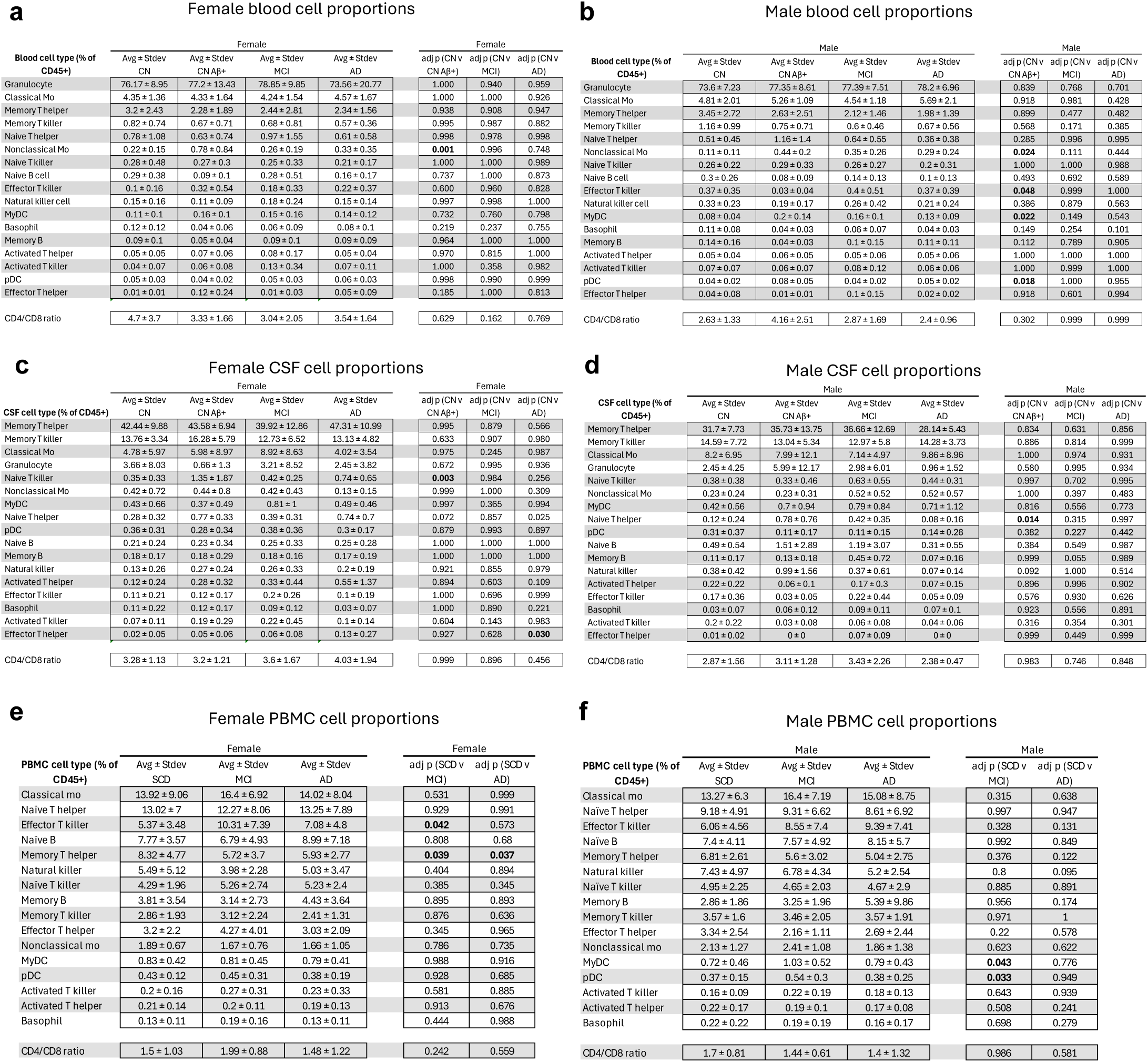
Cell type abundances across disease stages. **a,** Proportion of cells in female blood. (**a-f**) Two-way ANOVA with Dunnett multiple comparison of z-score normalized population abundance. **b**, Proportion of cells in male blood. **c**, Proportion of cells in female CSF. **d,** Proportion of cells in male CSF. **e,** Proportion of cells in female PBMC fraction. **f,** Proportion of cells in male PBMC fraction.

**Table 4.**
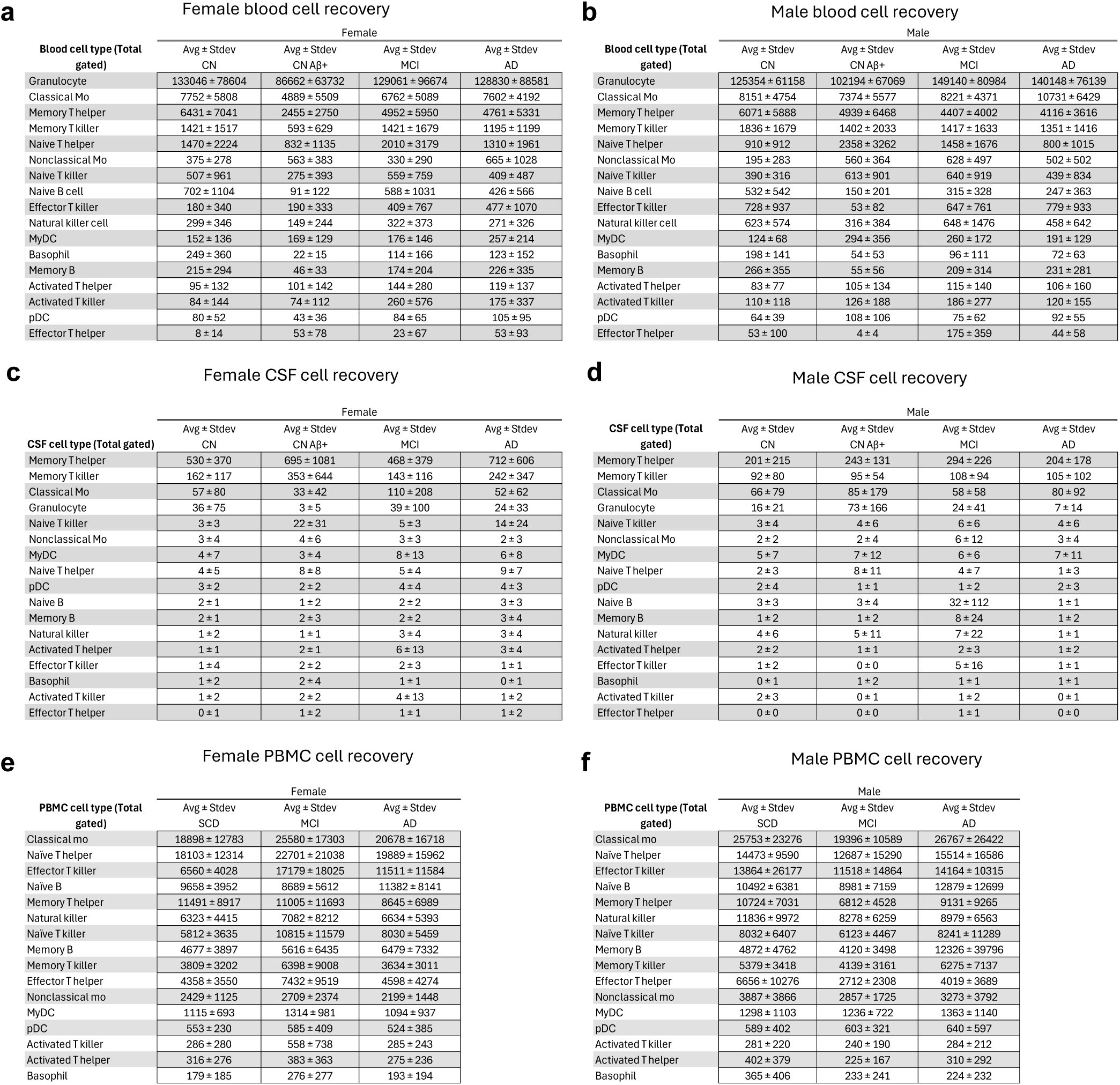
Cell type recovery across disease stages. **a**, Cells gated in female blood. **b**, Cells gated in male blood. **c**, Cells gated in female CSF. **d,** Cells gated in male CSF. **e,** Cells gated in female PBMC fraction. **f,** Cells gated in male PBMC fraction.

| Blood cell type (Total gated) | Female |  |  |  |
| --- | --- | --- | --- | --- |
|  | Avg + Stdev<br>CN | Avg + Stdev<br>CN Aβ+ | Avg + Stdev<br>MCI | Avg + Stdev<br>AD |
| Granulocyte | 133046 ± 78604 | 86662 ± 63732 | 129061 ± 96674 | 128830 ± 88581 |
| Classical Mo | 7752 ± 5808 | 4889 ± 5509 | 6762 ± 5089 | 7602 ± 4192 |
| Memory T helper | 6431 ± 7041 | 2455 ± 2750 | 4952 ± 5950 | 4761 ± 5331 |
| Memory T killer | 1421 ± 1517 | 593 ± 629 | 1421 ± 1679 | 1195 ± 1199 |
| Naive T helper | 1470 ± 2224 | 832 ± 1135 | 2010 ± 3179 | 1310 ± 1961 |
| Nonclassical Mo | 375 ± 278 | 563 ± 383 | 330 ± 290 | 665 ± 1028 |
| Naive T killer | 507 ± 961 | 275 ± 393 | 559 ± 759 | 409 ± 487 |
| Naive B cell | 702 ± 1104 | 91 ± 122 | 588 ± 1031 | 426 ± 566 |
| Effector T killer | 180 ± 340 | 190 ± 333 | 409 ± 767 | 477 ± 1070 |
| Natural killer cell | 299 ± 346 | 149 ± 244 | 322 ± 373 | 271 ± 326 |
| MyDC | 152 ± 136 | 169 ± 129 | 176 ± 146 | 257 ± 214 |
| Basophil | 249 ± 360 | 22 ± 15 | 114 ± 166 | 123 ± 152 |
| Memory B | 215 ± 294 | 46 ± 33 | 174 ± 204 | 226 ± 335 |
| Activated T helper | 95 ± 132 | 101 ± 142 | 144 ± 280 | 119 ± 137 |
| Activated T killer | 84 ± 144 | 74 ± 112 | 260 ± 576 | 175 ± 337 |
| pDC | 80 ± 52 | 43 ± 36 | 84 ± 65 | 105 ± 95 |
| Effector T helper | 8 ± 14 | 53 ± 78 | 23 ± 67 | 53 ± 93 |

**b** Male blood cell recovery
| Blood cell type (Total gated) | Male |  |  |  |
| --- | --- | --- | --- | --- |
|  | Avg + Stdev<br>CN | Avg + Stdev<br>CN Aβ+ | Avg + Stdev<br>MCI | Avg + Stdev<br>AD |
| Granulocyte | 125354 ± 61158 | 102194 ± 67069 | 149140 ± 80984 | 140148 ± 76139 |
| Classical Mo | 8151 ± 4754 | 7374 ± 5577 | 8221 ± 4371 | 10731 ± 6429 |
| Memory T helper | 6071 ± 5888 | 4939 ± 6468 | 4407 ± 4002 | 4116 ± 3616 |
| Memory T killer | 1836 ± 1679 | 1402 ± 2033 | 1417 ± 1633 | 1351 ± 1416 |
| Naive T helper | 910 ± 912 | 2358 ± 3262 | 1458 ± 1676 | 800 ± 1015 |
| Nonclassical Mo | 195 ± 283 | 560 ± 364 | 628 ± 497 | 502 ± 502 |
| Naive T killer | 390 ± 316 | 613 ± 901 | 640 ± 919 | 439 ± 834 |
| Naive B cell | 532 ± 542 | 150 ± 201 | 315 ± 328 | 247 ± 363 |
| Effector T killer | 728 ± 937 | 53 ± 82 | 647 ± 761 | 779 ± 933 |
| Natural killer cell | 623 ± 574 | 316 ± 384 | 648 ± 1476 | 458 ± 642 |
| MyDC | 124 ± 68 | 294 ± 356 | 260 ± 172 | 191 ± 129 |
| Basophil | 198 ± 141 | 54 ± 53 | 96 ± 111 | 72 ± 63 |
| Memory B | 266 ± 355 | 55 ± 56 | 209 ± 314 | 231 ± 281 |
| Activated T helper | 83 ± 77 | 105 ± 134 | 115 ± 140 | 106 ± 160 |
| Activated T killer | 110 ± 118 | 126 ± 188 | 186 ± 277 | 120 ± 155 |
| pDC | 64 ± 39 | 108 ± 106 | 75 ± 62 | 92 ± 55 |
| Effector T helper | 53 ± 100 | 4 ± 4 | 175 ± 359 | 44 ± 58 |

**c** Female CSF cell recovery
| CSF cell type (Total gated) | Female |  |  |  |
| --- | --- | --- | --- | --- |
|  | Avg + Stdev<br>CN | Avg + Stdev<br>CN Aβ+ | Avg + Stdev<br>MCI | Avg + Stdev<br>AD |
| Memory T helper | 530 ± 370 | 695 ± 1081 | 468 ± 379 | 712 ± 606 |
| Memory T killer | 162 ± 117 | 353 ± 644 | 143 ± 116 | 242 ± 347 |
| Classical Mo | 57 ± 80 | 33 ± 42 | 110 ± 208 | 52 ± 62 |
| Granulocyte | 36 ± 75 | 3 ± 5 | 39 ± 100 | 24 ± 33 |
| Naive T killer | 3 ± 3 | 22 ± 31 | 5 ± 3 | 14 ± 24 |
| Nonclassical Mo | 3 ± 4 | 4 ± 6 | 3 ± 3 | 2 ± 3 |
| MyDC | 4 ± 7 | 3 ± 4 | 8 ± 13 | 6 ± 8 |
| Naive T helper | 4 ± 5 | 8 ± 8 | 5 ± 4 | 9 ± 7 |
| pDC | 3 ± 2 | 2 ± 2 | 4 ± 4 | 4 ± 3 |
| Naive B | 2 ± 1 | 1 ± 2 | 2 ± 2 | 3 ± 3 |
| Memory B | 2 ± 1 | 2 ± 3 | 2 ± 2 | 3 ± 4 |
| Natural killer | 1 ± 2 | 1 ± 1 | 3 ± 4 | 3 ± 4 |
| Activated T helper | 1 ± 1 | 2 ± 1 | 6 ± 13 | 3 ± 4 |
| Effector T killer | 1 ± 4 | 2 ± 2 | 2 ± 3 | 1 ± 1 |
| Basophil | 1 ± 2 | 2 ± 4 | 1 ± 1 | 0 ± 1 |
| Activated T killer | 1 ± 2 | 2 ± 2 | 4 ± 13 | 1 ± 2 |
| Effector T helper | 0 ± 1 | 1 ± 2 | 1 ± 1 | 1 ± 2 |

**d** Male CSF cell recovery
| CSF cell type (Total gated) | Male |  |  |  |
| --- | --- | --- | --- | --- |
|  | Avg + Stdev<br>CN | Avg + Stdev<br>CN Aβ+ | Avg + Stdev<br>MCI | Avg + Stdev<br>AD |
| Memory T helper | 201 ± 215 | 243 ± 131 | 294 ± 226 | 204 ± 178 |
| Memory T killer | 92 ± 80 | 95 ± 54 | 108 ± 94 | 105 ± 102 |
| Classical Mo | 66 ± 79 | 85 ± 179 | 58 ± 58 | 80 ± 92 |
| Granulocyte | 16 ± 21 | 73 ± 166 | 24 ± 41 | 7 ± 14 |
| Naive T killer | 3 ± 4 | 4 ± 6 | 6 ± 6 | 4 ± 6 |
| Nonclassical Mo | 2 ± 2 | 2 ± 4 | 6 ± 12 | 3 ± 4 |
| MyDC | 5 ± 7 | 7 ± 12 | 6 ± 6 | 7 ± 11 |
| Naive T helper | 2 ± 3 | 8 ± 11 | 4 ± 7 | 1 ± 3 |
| pDC | 2 ± 4 | 1 ± 1 | 1 ± 2 | 2 ± 3 |
| Naive B | 3 ± 3 | 3 ± 4 | 32 ± 112 | 1 ± 1 |
| Memory B | 1 ± 2 | 1 ± 2 | 8 ± 24 | 1 ± 2 |
| Natural killer | 4 ± 6 | 5 ± 11 | 7 ± 22 | 1 ± 1 |
| Activated T helper | 2 ± 2 | 1 ± 1 | 2 ± 3 | 1 ± 2 |
| Effector T killer | 1 ± 2 | 0 ± 0 | 5 ± 16 | 1 ± 1 |
| Basophil | 0 ± 1 | 1 ± 2 | 1 ± 1 | 1 ± 1 |
| Activated T killer | 2 ± 3 | 0 ± 1 | 1 ± 2 | 0 ± 1 |
| Effector T helper | 0 ± 0 | 0 ± 0 | 1 ± 1 | 0 ± 0 |

**e** Female PBMC cell recovery
| PBMC cell type (Total gated) | Female |  |  |
| --- | --- | --- | --- |
|  | Avg + Stdev<br>SCD | Avg + Stdev<br>MCI | Avg + Stdev<br>AD |
| Classical mo | 18898 ± 12783 | 25580 ± 17303 | 20678 ± 16718 |
| Naive T helper | 18103 ± 12314 | 22701 ± 21038 | 19889 ± 15962 |
| Effector T killer | 6560 ± 4028 | 17179 ± 18025 | 11511 ± 11584 |
| Naive B | 9658 ± 3952 | 8689 ± 5612 | 11382 ± 8141 |
| Memory T helper | 11491 ± 8917 | 11005 ± 11693 | 8645 ± 6989 |
| Natural killer | 6323 ± 4415 | 7082 ± 8212 | 6634 ± 5393 |
| Naive T killer | 5812 ± 3635 | 10815 ± 11579 | 8030 ± 5459 |
| Memory B | 4677 ± 3897 | 5616 ± 6435 | 6479 ± 7332 |
| Memory T killer | 3809 ± 3202 | 6398 ± 9008 | 3634 ± 3011 |
| Effector T helper | 4358 ± 3550 | 7432 ± 9519 | 4598 ± 4274 |
| Nonclassical mo | 2429 ± 1125 | 2709 ± 2374 | 2199 ± 1448 |
| MyDC | 1115 ± 693 | 1314 ± 981 | 1094 ± 937 |
| pDC | 553 ± 230 | 585 ± 409 | 524 ± 385 |
| Activated T killer | 286 ± 280 | 558 ± 738 | 285 ± 243 |
| Activated T helper | 316 ± 276 | 383 ± 363 | 275 ± 236 |
| Basophil | 179 ± 185 | 276 ± 277 | 193 ± 194 |

**Table 4** | Cell type recovery across disease stages.
| PBMC cell type (Total gated) | Male |  |  |
| --- | --- | --- | --- |
|  | Avg + Stdev<br>SCD | Avg + Stdev<br>MCI | Avg + Stdev<br>AD |
| Classical mo | 25753 ± 23276 | 19396 ± 10589 | 26767 ± 26422 |
| Naive T helper | 14473 ± 9590 | 12687 ± 15290 | 15514 ± 16586 |
| Effector T killer | 13864 ± 26177 | 11518 ± 14864 | 14164 ± 10315 |
| Naive B | 10492 ± 6381 | 8981 ± 7159 | 12879 ± 12699 |
| Memory T helper | 10724 ± 7031 | 6812 ± 4528 | 9131 ± 9265 |
| Natural killer | 11836 ± 9972 | 8278 ± 6259 | 8979 ± 6563 |
| Naive T killer | 8032 ± 6407 | 6123 ± 4467 | 8241 ± 11289 |
| Memory B | 4872 ± 4762 | 4120 ± 3498 | 12326 ± 39796 |
| Memory T killer | 5379 ± 3418 | 4139 ± 3161 | 6275 ± 7137 |
| Effector T helper | 6656 ± 10276 | 2712 ± 2308 | 4019 ± 3689 |
| Nonclassical mo | 3887 ± 3866 | 2857 ± 1725 | 3273 ± 3792 |
| MyDC | 1298 ± 1103 | 1236 ± 722 | 1363 ± 1140 |
| pDC | 589 ± 402 | 603 ± 321 | 640 ± 597 |
| Activated T killer | 281 ± 220 | 240 ± 190 | 284 ± 212 |
| Activated T helper | 402 ± 379 | 225 ± 167 | 310 ± 292 |
| Basophil | 365 ± 406 | 233 ± 241 | 224 ± 232 |

### Alterations in cellular abundance across clinical stages of Alzheimer’s disease

To determine whether the abundance of major immune populations was altered in people with CNS pathology related to AD, we compared the frequency of cell types in each stage relative to CN individuals or SCD controls using a two-way ANOVA with Dunnett’s correction for multiple comparisons. Within the innate immune compartment, nonclassical monocytes were enriched in the blood of female and male CN Aβ+ individuals (**Table 3**). We also found enrichment of myeloid dendritic cells (myDC) and plasmacytoid dendritic cells (pDC) in the blood of CN Aβ+ males and in the PBMC fraction of males with MCI (**Table 3**). Within the adaptive immune compartment, naï ve T helper cells were enriched in the CSF of male CN Aβ+ and effector T helper cells were enriched in the CSF of female AD individuals (**Table 3**). The only depleted population we observed was effector T killer cells, which were reduced in the blood of male CN Aβ+ individuals (**Table 3**).

Since CD8+ T_EMRA_ cells were reported to be elevated in PBMC from MCI and AD individuals in the second cohort^13^, we next investigated how the intensity of CD45RA, a defining marker of T_EMRA_ cells, varied across AD stages within the discovery cohort’s blood and CSF data (**Extended data Fig 2a-d**). CD45RA was elevated in circulating naï ve T killer cells (p-value = 0.008; two-way ANOVA with Dunnett’s correction) and naï ve T helper cells (p-value = 0.012) of CN Aβ+ females, as well as naï ve T helper cells (p-value = 0.047) of MCI females (**Extended data Fig 2a**). No significant CD45RA changes were observed in a sex or stage-specific manner among other T cell subsets in blood or CSF (**Extended data Fig 2a-d**). Together, our findings are consistent with the reported accumulation of peripheral monocytes, DCs and T_EMRA_ cells in conjunction with β amyloid deposition in the CNS^7–13^ and suggest that these innate and adaptive immune cell alterations in the periphery and CNS are both diminished in individuals with cognitive impairment related to AD.

### Sex-related influence of the immune proteome

Sex is a major modifier of transcriptional programs in microglia and peripheral immune cells isolated from cognitively normal adults and those with AD neuropathology^11,32–34^. To examine whether sex affected the proteome of immune populations in the periphery and CNS, we compared the average intensity of immune features in men and women classified by their AD disease stage. We observed an inverse relationship between the average intensity of immune features in male relative to female cells derived from blood, CSF and PBMC (**Extended data Fig 3a)**, indicating that the protein abundance for features we measured varied in opposite directions between the sexes. Next, to pinpoint immune features that were most affected by sex, we performed a linear regression analysis. Following Benjamini-Hochberg correction for multiple testing, we found seven features that were enriched in female circulating immune cells and ten enriched in male circulating immune cells, whereas no immune features from the CSF correlated with sex (**Extended data Fig 3b-d**). Notably, proteins associated with cytotoxicity, which included CD56 and CD57^35,36^ were enriched in male natural killer cells (NK) (**Extended data Fig 3e**), consistent with prior studies showing NK cells derived from adult males express higher levels of transcripts related to cytotoxicity but are functionally impaired relative to NK cells isolated from adult females^34,37^. Additionally, we found an enrichment of CD11c on circulating NK cells in females, an integrin involved in promoting complement-mediated phagocytosis^38^, as well as enrichment of the co-stimulatory molecule CD27 on female myDC that potentiates cytotoxic T cell responses to foreign antigens^39^ (**Extended data Fig 3e-f**).

To investigate how sex-dependent immune signatures varied across individuals and whether immune signatures were unique or shared across AD stages, we performed a principal component analysis (PCA) using sex-enriched blood features as inputs. The first principal component (21.7% of the variance) clearly separated males from females that were classified as CN, MCI or AD dementia. A similar trend was observed between male and female CN Aβ+ individuals (**Extended data Fig 3g**). When sex-enriched PBMC features were used as inputs, the first principal component (37.0% of the variance) separated males and females with SCD and AD and a trend was apparent to segregate MCI individuals by sex (**Extended data Fig 3h**). These findings underscore that sex is an important governor of the homeostatic set point of both the peripheral and CNS immune response and that sex-dependent differences in protein abundance persist in immune cells derived from individuals with AD pathology.

### Disease stage-enriched immune features

We next aimed to identify disease stage specific immune factors by comparing the abundance of immune features in CSF, blood and PBMC in a pairwise manner across AD stages using two-way ANOVA with Tukey correction for multiple comparisons (**Fig 1b**). In total, we identified 141 differentially abundant features in females (37 CSF features; 93 blood features, 11 PBMC features) and 101 differentially abundant features in males (28 CSF features; 66 blood features; 7 PBMC features) that met the adjusted threshold of p-value < 0.05 (**Fig 1c**). The larger number of differentially abundant immune features we observed in blood could be due to higher measurement precision, however true biological differences between compartments may also contribute to these effects (**Extended data Fig 1e-f** and **Table 4**). Immune features in the periphery exhibited stronger differences than those observed in the CSF.

Considering the class of immune cell types that participate in AD stage-related immune alteration, we found that females exhibited a greater number of differentially abundant innate immune features (27 CSF features; 53 blood features; 6 PBMC features) than adaptive immune features (10 CSF features; 40 blood features; 5 adaptive PBMC features) (**Fig 1d-f**). Conversely, we found that males displayed a greater number of differential abundant adaptive immune features (22 CSF features; 33 blood features; 6 PBMC features) than innate immune features (7 CSF features; 33 blood features; 1 PBMC feature) (**Fig 1d-f**). NK features were prominently represented in female CSF datasets (**Fig 1g**), whereas classical and nonclassical monocytic features were strongly represented in female blood (**Fig 1h**) and PBMC datasets (**Fig 1i**). Features from naive B cells were well represented in male CSF (**Fig 1g**), blood (**Fig 1h**) and PBMC datasets (**Fig 1i)**. The transition from CN to CN Aβ+ biological stage was marked by activation of naï ve T killer cells and suppression of nonclassical monocytes in blood of females and males (**Extended data Fig 4a-b**). Progression from CN Aβ+ to MCI was marked by suppression of circulating pDC activity in both sexes, along with recovery of nonclassical monocyte activation state in females with MCI (**Extended data Fig 4c-d**). Reduced T killer activity was evident in the MCI stage of males and the AD dementia stage in females (**Extended data Fig 4c-f**). Overall, biological sex appears to influence both the timing and nature of immune proteomic alterations associated with AD neuropathology.

### Stratification of individuals using immune profiles

Although this study was not powered to evaluate the diagnostic performance of CyTOF-based immune profiling, we performed an exploratory assessment on whether immune features derived from CyTOF capture biologically meaningful structure across the AD continuum and support clinical and biological stage-associated differences. To this end, we applied Andrew’s curves^40^, a dimensionality reduction approach that transforms multivariate measurements into a single function per individual, using blood-derived immune features. Andrew’s curves generated from immune features exhibited qualitative stratification by AD clinical stage that was comparable to that observed using conventional clinical and biofluid measures, including the Clinical Dementia Rating scale and established AD biomarkers (**Extended Data Fig. 5a–h**).

In parallel, we performed PCA using AD stage-enriched immune features from CSF (**Extended data Fig 6a-f**), blood (**Extended data figure 7a-f**) and PBMC (**Extended data Fig 8a-f**) as inputs. Across both compartments, separation of individuals by AD clinical stage was observed along the first or second principal component, with group differences assessed using one-way ANOVA with Sidak’s multiple-comparison test or Kruskal–Wallis with Dunn’s multiple-comparison test, as appropriate. These analyses illustrate the presence of coordinated, stage-associated immune variation and support future studies formally evaluating the utility of immune profiling for subgroup identification and biomarker development.

### Altered immune modules in the CNS across AD biological and clinical stages

Given the high dimensionality of CyTOF data, we defined co-aggregated immune modules to summarize correlated features into biologically coherent immune programs that could be compared across compartments, AD stages, and cohorts. Our approach was based on principles of weighted gene co-expression network analysis^41^ in which features with similar expression patterns are predicted to be functionally associated due to physical interactions, involvement in shared biological pathways, or regulation by common mechanisms. We constructed a covariance matrix using CSF-derived immune features and applied a threshold to enrich for highly correlated features. This analysis revealed two modules from the CSF of females (modules 1 and 2) (**Fig 2a-d**) and from the CSF of males (modules 3 and 4) (**Fig 2e-h**).

**Fig. 2.**
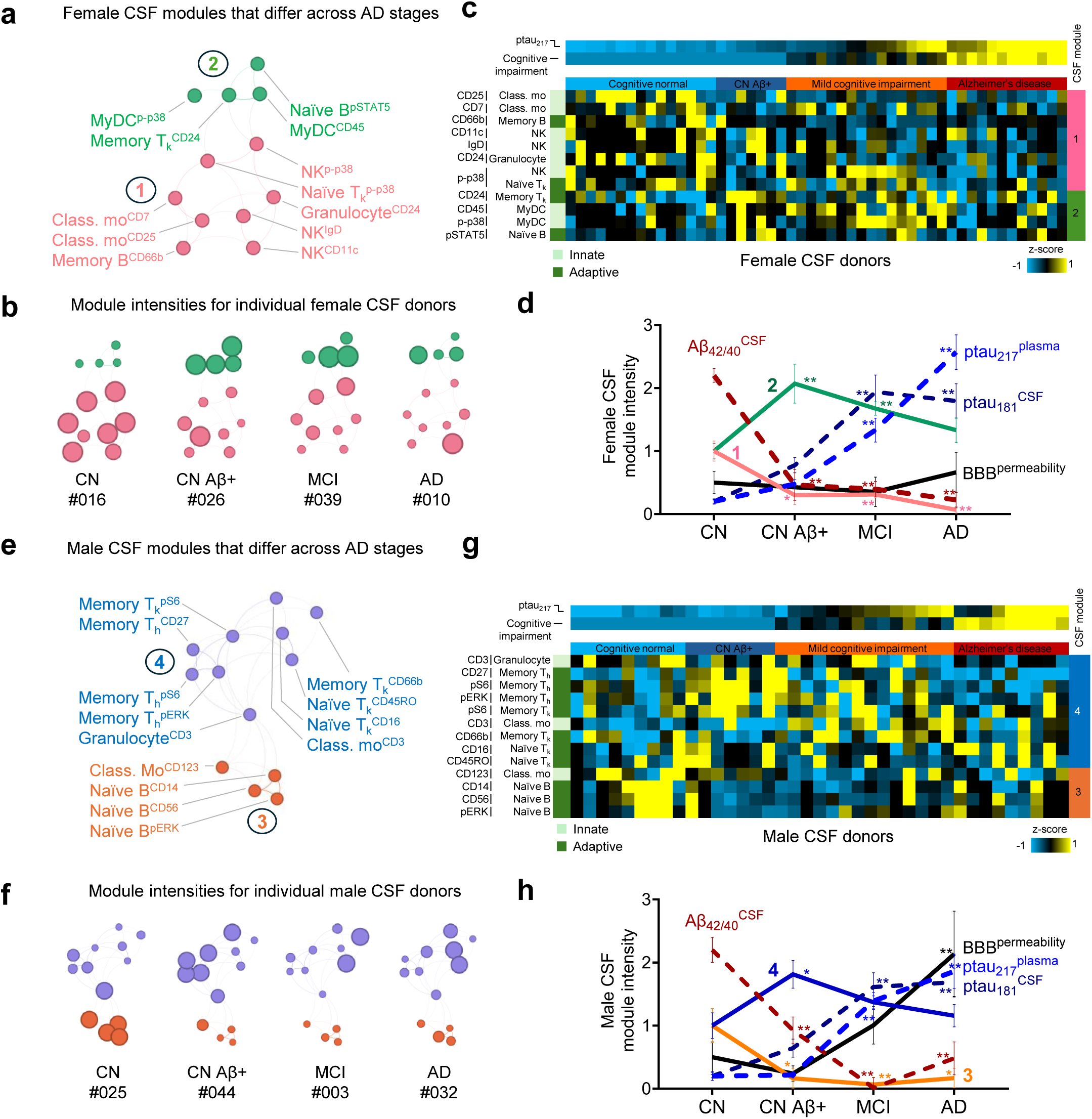
Dynamic sex-specific central nervous system immune networks across clinical stages of Alzheimer’s disease. **a-h,** Networks of female (**a-d**) or male (**e-h**) CSF immune features colored by module with connections representing the strength of feature covariance (**a, b, e, f**). **b, f**, Network node sizes were scaled by feature intensity for representative CSF donors classified by disease stage and patient identifier number. **c, g,** The intensity of features within female (**c**) or male (**g**) CSF donors were colored by z-score normalized intensity (teal low; yellow high). Plasma ptau_217_ and cognitive impairment represented by Clinical dementia rating sum of boxes are reported above for each donor. **d, h**, Summary of CSF module dynamics across disease stages. Hippocampal blood brain barrier (BBB) permeability was gleaned from DCE-MRI. Error bars represent S.E.M. Two-way ANOVA with Dunnett’s multiple comparison test relative to CN. * p < 0.05 ** p < 0.01. Activated (Act.). Effector (Eff.). Memory (Mem.). CD8+ T killer cell (T_k_). CD4+ T helper cell (T_h_). Myeloid dendritic cell (myDC). Plasmacytoid dendritic cell (pDC). Classical monocyte (Class. mo). Nonclassical monocyte (Nonclas. mo). Natural killer cell (NK).

To visualize how immune features varied in relation to one another, we represented each feature – the combination of cell type and protein - as a node within a network colored by module class, with edges weighted by the degree of covariance between the nodes (**Fig 2a** and **Fig 2e**). We then applied unsupervised hierarchical clustering to arrange the immune features within a heat map and ordered donor samples based on the severity of clinical impairment as measured by the Clinical Dementia Rating sum of boxes^42^ and levels of plasma phospho-tau 217 (ptau_217_), an early indicator of AD neuropathological changes^43^ (**Fig 2c** and **Fig 2g**). Finally, we applied a two-way ANOVA with Dunnett’s correction to compare the intensity of CSF immune modules in each disease stage. From simultaneously isolated samples, we compared the Amyloid β_42/40_ ratio in CSF that is indicative of Aβ deposition in the CNS, as well as measures of plasma ptau_217_, CSF ptau_181_, and hippocampal BBB permeability (**Fig 2d** and **Fig 2h**).

Female CSF module 1, reduced in CN Aβ+, MCI and AD stages relative to CN individuals, was characterized by expression of the CD7 glycoprotein and the high affinity IL-2 receptor CD25 on classical monocytes as well as the p-p38 signaling molecule within NK and naï ve T killer cells (**Fig 2d**). CSF module 2, enriched in CN Aβ+ and MCI females but not altered in AD females, was defined by accumulation of p-p38 and CD45 in myDC as well as the checkpoint inhibitor CD24^44^ on memory T killer cells (**Fig 2d**). A similar pattern of proteomic immune alterations was observed in CSF cells derived from males. Module 3 was defined by naï ve B cell expression of the LPS co-receptor CD14, CD56 and pErk signaling molecule and was reduced in CN Aβ+, MCI and AD stages relative to CN individuals (**Fig 2h**). Module 4 was elevated uniquely in CN Aβ+ males and was enriched for the cell adhesion molecule CD66b on memory T killer cells as well as the T cell co-receptor CD3 on granulocytes and classical monocytes (**Fig 2h**). The presence of granulocyte-restricted CD66b on memory T killer cells and CD3 on granulocytes and monocytes is suggestive of trogocytosis^45^, a process in which membrane proteins are transferred from one cell to another following physical contact. The findings suggest that in the local CNS microenvironment of CN Aβ+ males, the presence of β amyloid aggregates may trigger granulocytes to act as antigen presenting cells that engage T killer cells.

We also examined whether carriage of the *APOE ε4* risk allele was associated with differences in CSF immune module intensity across the AD continuum. We did not observe a significant effect of *APOE ε4* status on the relative abundance of CSF immune modules, as ε4 carriers and non-carriers showed comparable module intensities across AD stages when assessed by two-way ANOVA with Sidak’s multiple-comparisons test (all adjusted p values > 0.1; **Extended Data Fig. 9a-b**).

Importantly, we found that immune alterations occurred in the CSF of CN Aβ+ individuals in the absence of significant BBB impairment or elevated plasma ptau_217_ or CSF ptau_181_. In contrast, phosphorylated tau levels were notably elevated in the MCI and AD stage of disease, whereas hippocampal BBB permeability was primarily elevated in males with AD dementia (**Fig 2d** and **Fig 2h**). Together, these findings indicate that CSF immune state changes are detectable even during the preclinical stage of AD and occur prior to widespread tau pathology and overt BBB disruption as measured by sensitive MRI-based assessments of molecular permeability.

### Peripheral immune modules vary across AD biological and clinical stages

Next, we applied a similar strategy to identify highly correlated biologically coherent peripheral immune modules that varied across AD stages. We recovered five modules from the blood of females (modules 5-9) (**Fig 3a-d**) and four modules from the blood of males (modules 10-13) (**Fig 3e-h**). Modules 7, 8 and 11 were enriched in CN Aβ+ individuals whereas modules 6 and 10 were reduced in CN Aβ+ individuals relative to CN controls (**Fig 3d** and **Fig 3h**). CD33, highly enriched in module 7, functions as a receptor for sialic acid that influences microglial and monocyte phagocytic activity^46^ and is implicated as a risk factor for late onset AD^47–49^. Module 8 featured granulocyte expression of phospho-pPLCγ2 and phospho-STAT5, the IL-7 receptor CD127 and CD8, along with pAkt, pSTAT3 and pSTAT5 in naï ve T killer cells, suggesting cytokine receptor signaling driven coordination^50^. A bidirectional relationship between antigen presentation and signaling downstream of mTOR in CN Aβ+ males was suggested by accumulation of the module 11 components HLA-DR on pDC, naï ve B and activated T killer cells as well as the elevation of pS6 in pDC and activated T helper cells. The reduction of the IL-3 receptor CD123 on classical and nonclassical monocytes in CN Aβ+ females as components of module 6 (**Fig 3d**) may be a compensatory response to limit trained immunity and inflammation^51^. Intriguingly, module 6 was reduced in CN *APOE ε4* carriers compared to non-carriers (adjusted p-value < 0.05) (**Extended data Fig 10a**). Notably, CD123 was reduced on circulating naï ve B and memory B cells within module 10 of CN Aβ+ males (**Fig 3h**).

**Fig. 3.**
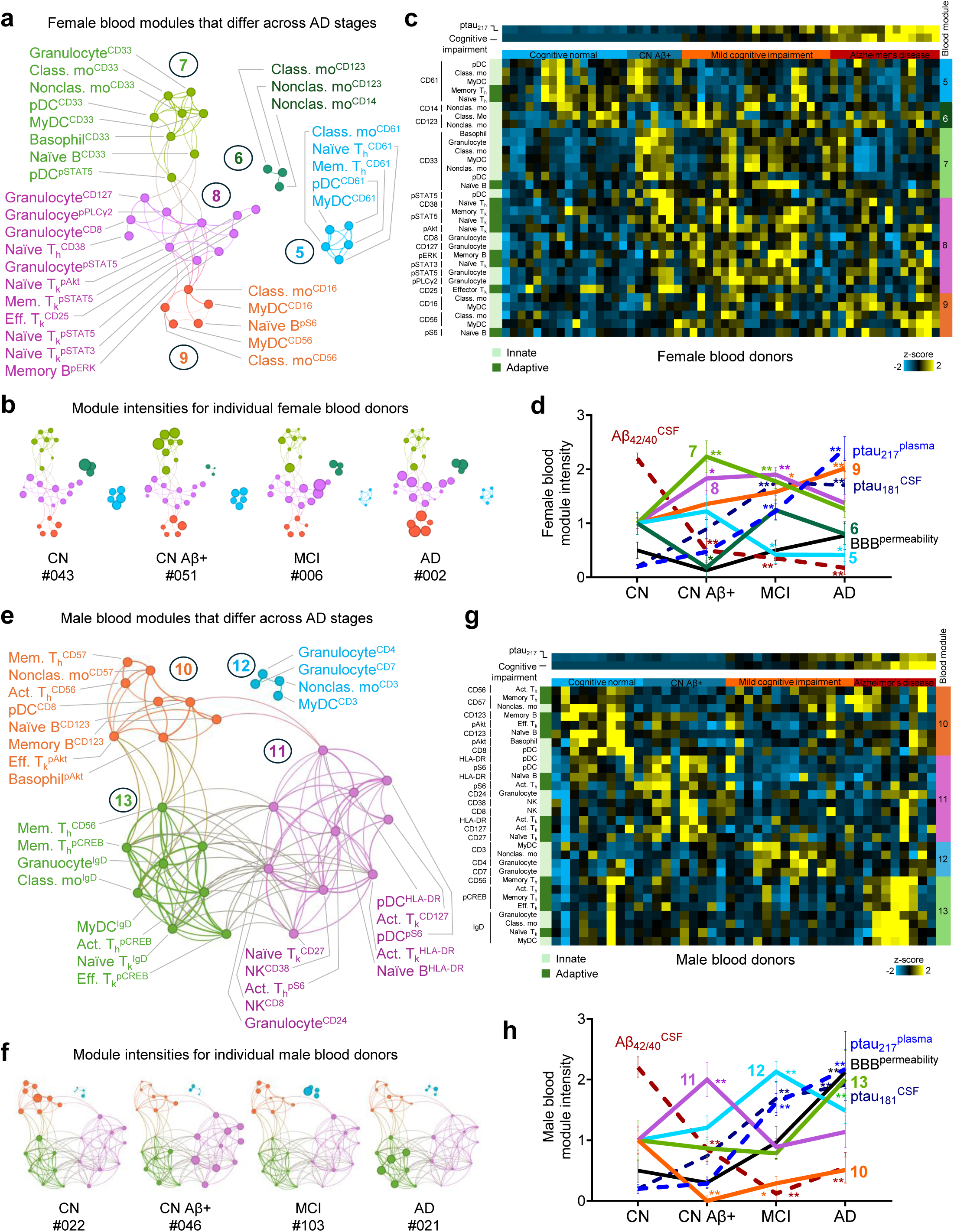
Dynamic sex-specific peripheral immune networks across clinical stages of Alzheimer’s disease. **a-h,** Network of female (**a-d**) or male (**e-h**) peripheral blood immune features colored by module with connections representing the strength of feature covariance (**a, b, e, f**). **b, f**, Network node sizes were scaled by marker intensity for representative individuals classified by disease stage and patient identifier number. **c**, **g**, Intensity of module components across female (**c**) or male (**g**) blood donors colored by z-score normalized intensity (teal low; yellow high). Plasma ptau_217_ and cognitive impairment represented by Clinical dementia rating sum of boxes are reported above. **d, h**, Summary of module dynamics across disease stages. Hippocampal blood brain barrier (BBB) permeability gleaned from DCE-MRI. Error bars represent S.E.M. Two-way ANOVA with Dunnett’s multiple comparison test relative to CN. * p < 0.05 ** p < 0.01. See Fig. 2 legend for list of abbreviations.

MCI stage of AD was associated with an enrichment of modules 7, 8 and 12 in peripheral blood and depletion of modules 5 and 10 (**Fig 3d** and **Fig 3h**). Module 12 captured the coordinated accumulation of CD3 on nonclassical monocytes and myDC and enrichment of CD4 on granulocytes (**Fig 3e**), suggesting interaction between these innate immune cells and T cells. The reduction of module 5, dominated by the β3 integrin CD61 that is important for binding fibrinogen to induce platelet aggregation^52^, may be indicative of increased consumption of clotting factors in females with MCI.

We found that AD dementia stage was uniquely associated with elevated modules 9 and 13 in peripheral blood, while modules 5 and 10 remained consistently reduced in AD dementia cases relative to CN controls (**Fig 3d** and **Fig 3h**). Module 9 featured enrichment of CD56 and the Fc gamma receptor III CD16 on myDC and classical monocytes, a phenotype that emulated proinflammatory cell states observed in patients with Rheumatoid arthritis^53^. Module 13 showed elevation of CD56 on memory T helper cells along with enrichment of pCREB within innate and adaptive immune cells coated with immunoglobulin IgD (**Fig 3e**). Although module 7 was not altered in AD dementia relative to CN controls, its CD33-features were elevated in female *APOE ε4* positive AD dementia individuals (p-value < 0.01)(**Extended data Fig 10a**). By aggregating immune features that correlated linearly with plasma ptau_217_ levels, we identified 6 additional sex-specific blood modules (female modules 14-16 and male modules 17-19) that varied across AD stages (**Extended data Fig 11a-j**). The finding that peripheral immune alterations in CN Aβ+ individuals precede the significant accumulation of plasma ptau_217_, CSF ptau_181_ and notable BBB permeability on DCE MRI (**Fig 3d** and **Fig 3h**) further supports the idea that systemic immune changes are an early response rather than a later consequence of increasing AD pathology burden.

### Signal transduction proteins drive AD stage-specific hubs of influence

We next sought to identify signaling proteins that were most important for coordinating immune responses in the periphery and CNS at each biological and clinical stage of AD. To do so, we evaluated the full landscape of immune features, including proteins that did not exhibit significant stage-associated differences, reasoning that regulators of immune network structure may exert influence through coordinated interactions rather than through large univariate changes. We stratified the CyTOF datasets by sex, disease state and tissue of origin and generated covariance networks that encompassed multiple immune features we had measured (**Fig 4a-b and Video S1**).

**Fig. 4.**
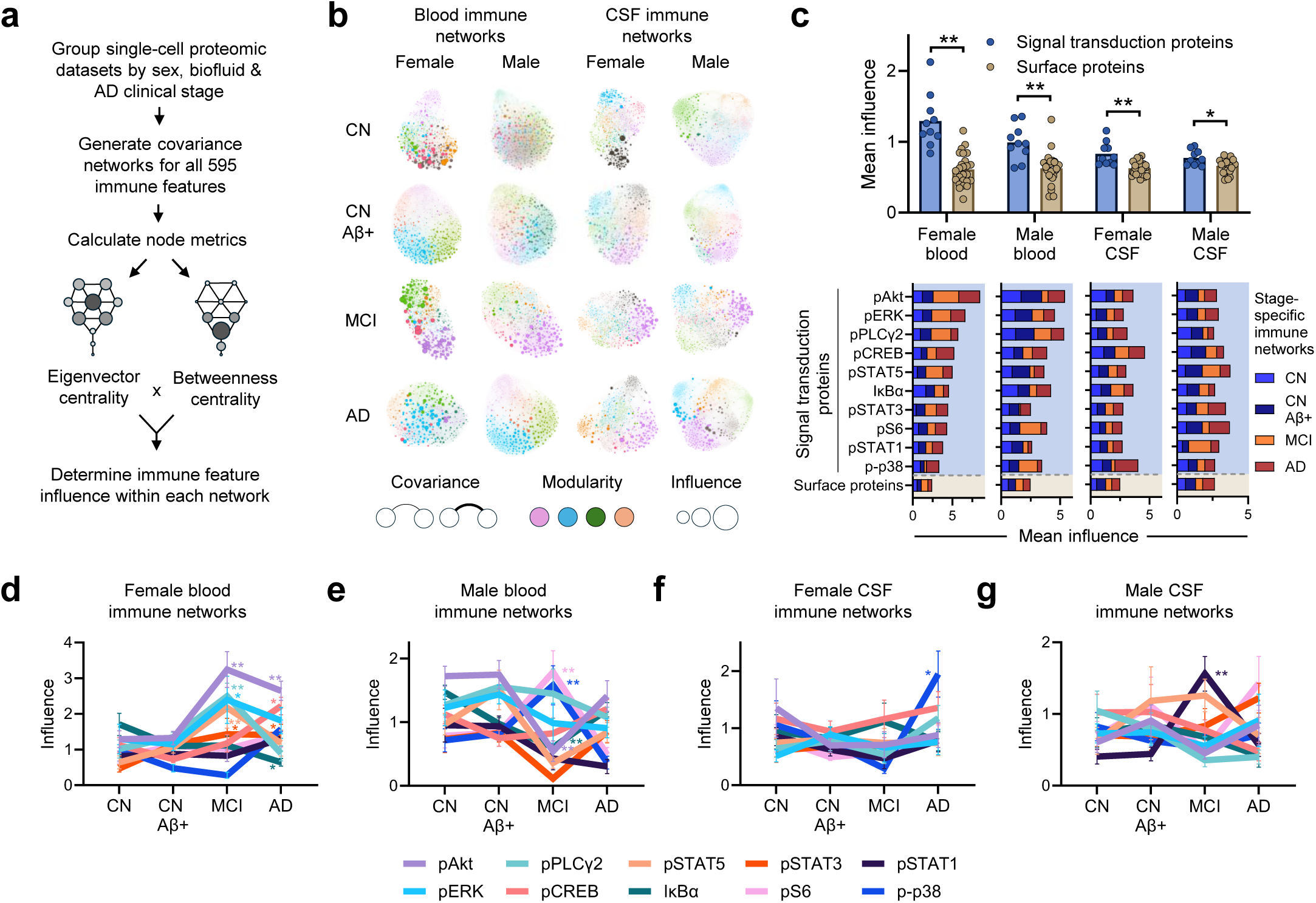
Signal transduction proteins coordinate peripheral and CNS immune activity across clinical stages of Alzheimer’s disease. **a,** Strategy to derive covariance networks of immune features stratified by sex, biofluid and AD clinical stage. **b**, Each image represents a network of 595 nodes comprising cell type – antigen combinations (features). Connections are the weighted strength of covariance between nodes. Node color relates to modularity. Node size is proportional to influence of each immune feature, derived by multiplying the value of betweenness centrality and eigenvector centrality within each network. See Video S1. **c,** Mean influence of signal transduction and surface protein features within each sex-stratified tissue-specific network (top) or broken out by disease stage-specific networks (below). Welch’s t-test for female blood, female CSF and male CSF. Mann-Whitney test for male blood. ** p < 0.01. **d-g**, Summary of signaling protein influence across disease stages for (**d**) female blood immune networks, (**e**) male blood immune networks, (**f**) female CSF immune networks and (g) male CSF immune networks). Two-way ANOVA with Dunnett’s multiple comparison test relative to CN. * p < 0.05 ** p < 0.01. Error bars represent SEM.

Within each network, immune features were represented as nodes colored by module class, with connection strengths weighted by feature covariance. To assess node importance, we calculated eigenvector centrality and betweenness centrality^54^. Nodes with high eigenvector centrality were directly connected to other highly connected nodes, whereas nodes with high betweenness centrality lie on the shortest path between other nodes in the network. Multiplying eigenvector centrality and betweenness centrality yielded an overall influence score for each node.

Strikingly, for both blood and CSF, we found that proteins involved in signal transduction were significantly more influential than surface receptors (**Fig 4c** and **Extended data Fig 12a-b**), suggesting that common signaling pathways are activated in a compartment specific manner to coordinate immune responses involving multiple cell types. Within the PBMC dataset, we identified highly influential features that associated with T cell costimulation that included CD5, TCRγδ, CTLA4, ICOS and TIGIT (**Extended data Fig 12c-d**). The predominance of signaling proteins as influential features likely reflects their ability to capture integrated cellular activation states downstream of multiple upstream related inputs (cytokines, antigens, metabolic pathways). In contrast, surface receptor expression is often more cell-type–restricted and context-dependent, which may limit its influence in network-based analyses. Although intracellular signaling proteins were not profiled in PBMC within the Amsterdam cohort, the most influential PBMC features were enriched for T cell costimulatory and inhibitory receptors that regulate downstream signaling pathways, again indicating consistency across cohorts and compartments.

We next examined how sex-specific immune cell activations varied across AD biological and clinical stages. We found no major difference in the relative influence of signaling proteins in blood or CSF in preclinical AD (**Fig 4d-g**). However, the influence of pERK, pPLCγ2, pAkt, and pSTAT3 was enriched in the blood of females with MCI (**Fig 4d**), with pAkt and pSTAT3 remaining persistently elevated in the blood of females with AD dementia (**Fig 4d**). By contrast, p-p38 and pS6 influence was enriched and IκBα and pAkt were reduced in the blood of males with MCI (**Fig 4e**). Within the CSF compartment of females with AD dementia, p-p38 signaling was most influential (**Fig 4f**), whereas pSTAT1 was influential in the CSF compartment of males with MCI (**Fig 4g**). Together, these results illustrate that while the earliest signaling protein activation in preclinical AD appears comparable between sexes, AD progression by disease stage is associated with sex and compartment specific reweighting of intracellular signaling pathways that coordinate immune responses differently between sexes.

### Elastic net identifies robust immune processes altered in AD

To prioritize immune features related to AD pathologic processes, we applied an orthogonal modeling strategy to perform variable selection across the entire set of blood CyTOF-based immune features identified in a univariate model involving baseline plasma ptau_217_ levels (84 features were included at p-value < 0.05). To address the extensive subject missingness within the effector T helper cell population, features from this cell type were consolidated into a binary trait that identified the presence or absence of detectable effector T helper cells (**Extended data Fig 1f**). We performed this analysis on males and females jointly (adjusting for *APOE* status and age) to optimize statistical power for the multivariable analysis and to identify the peripheral immune features that generalize best across sexes.

Elastic net regression^55^ on baseline plasma ptau_217_ levels identified 20 significant features. Importantly, the strongest predictor of increased plasma ptau_217_ (activated T helper^CD7^) and the strongest predictor of decreased plasma ptau_217_ (memory B^CD14^) were also identified in sex-stratified analyses, driven predominantly by effects in females. Of 20 features, eight either overlapped or were strongly correlated (corr > 0.5) with features identified in sex-stratified analyses. These features cluster loosely into three modules (**Extended data Fig 13a-f**). 1) An “intact cognition” module characterized by elevated memory B^CD14^ and naï ve B^CD61^ that correspond to lymphocyte activation^56^ with stronger effects in females (correlated to module 14) and that tended to be elevated in CN controls (**Extended data Fig 13a-b**). 2) An “AD progression” module typified by memory B^IgD^ and granulocyte^IκBα^, reflecting NF-κB-driven chronic adaptive^57^ and innate^58^ immune activation, which correlated to module 19, yet affecting both males and females, and that tended to become elevated with the extent of cognitive impairment related to AD (**Extended data Fig 13a-b**). 3) A “T cell exhaustion” module characterized by activated T helper^CD7^ and naï ve T helper^CD7^ that correspond to a shift in early-differentiated T helper cell populations^59,60^ and correlated to modules 16 and 18, but with somewhat stronger effects in males, and that tended to be enriched in subjects with AD dementia (**Extended data Fig 13a-b**).

Using ordinal elastic net regression, we identified peripheral immune features that track transitions across AD biological and clinical stages. As input, we used a set of significant features derived from univariate analysis (67 features were included at p-value < 0.05 and the effector T helper cell population was binary (detected/not detected)). Our ordinal elastic net identified predictors (either directly or through strong correlation) from four of the six blood modules defined within females, and all blood modules within males. Primary features that drive these modules include memory B^CD123^ (module 10), myDC^CD16^ (module 9), memory T killer^CD56^ (module 13), basophil^CD19^, myDC^CD123^, naï ve B^CD33^ (module 7), and granulocyte^pPLCγ2^ (module 8) (**Extended data Fig 13c-d**). For females, only module six (monocytic features depleted in CN Aβ+ stage) did not have a representative or correlated feature selected in our sex-combined elastic net analyses. Our ordinal prediction model shows strong agreement with disease classification (Cohen’s Kappa = 0.76 [0.64-0.88]), though notably the model selectively classifies participants into two groups, CN vs. CN Aβ+/MCI/AD.

The dichotomization of individuals into one “case” group by the model implies that peripheral immune system changes reflect underlying AD pathology and its biological stage more closely than clinical stages of AD. These multivariable elastic net analyses reduce the influence of correlated features to identify memory B^CD14^, naï ve B^CD61^, activated T helper^CD7^, memory B^CD123^, and myDC^CD16^ as the strongest peripheral immune features that track plasma ptau_217_ levels, and AD status, highlighting candidate early immune signatures that distinguish CN individuals without amyloid from those with preclinical or clinical stages of AD.

### Soluble factors promote AD-specific immune cell signatures

Lastly, we sought to gain insight into the biological drivers of immune cell activation signatures we report and evaluate their specificity with relation to underlying AD neuropathology. We exposed freshly isolated blood cells from three independent CN donors to plasma or CSF derived from either CN control or AD dementia females followed by mass cytometry evaluation (**Fig 5a**). Each stimulatory condition engaged a largely unique set of differentially abundant immune features when compared with vehicle treated control cells (**Fig 5b**), which could be further stratified into modular networks of covariable immune features (**Fig 5c**). Exposure of CN blood cells to AD plasma or CSF, but not CN plasma or CSF, induced specific immune cell signaling events that mirror the immune features identified in our MCI and AD patient cohorts (**Fig 5d**). These included elevated pAkt in naï ve T killer cells (**Fig 5e**) and increased pPLCγ2 in granulocytes (**Fig 5f**).

**Fig. 5.**
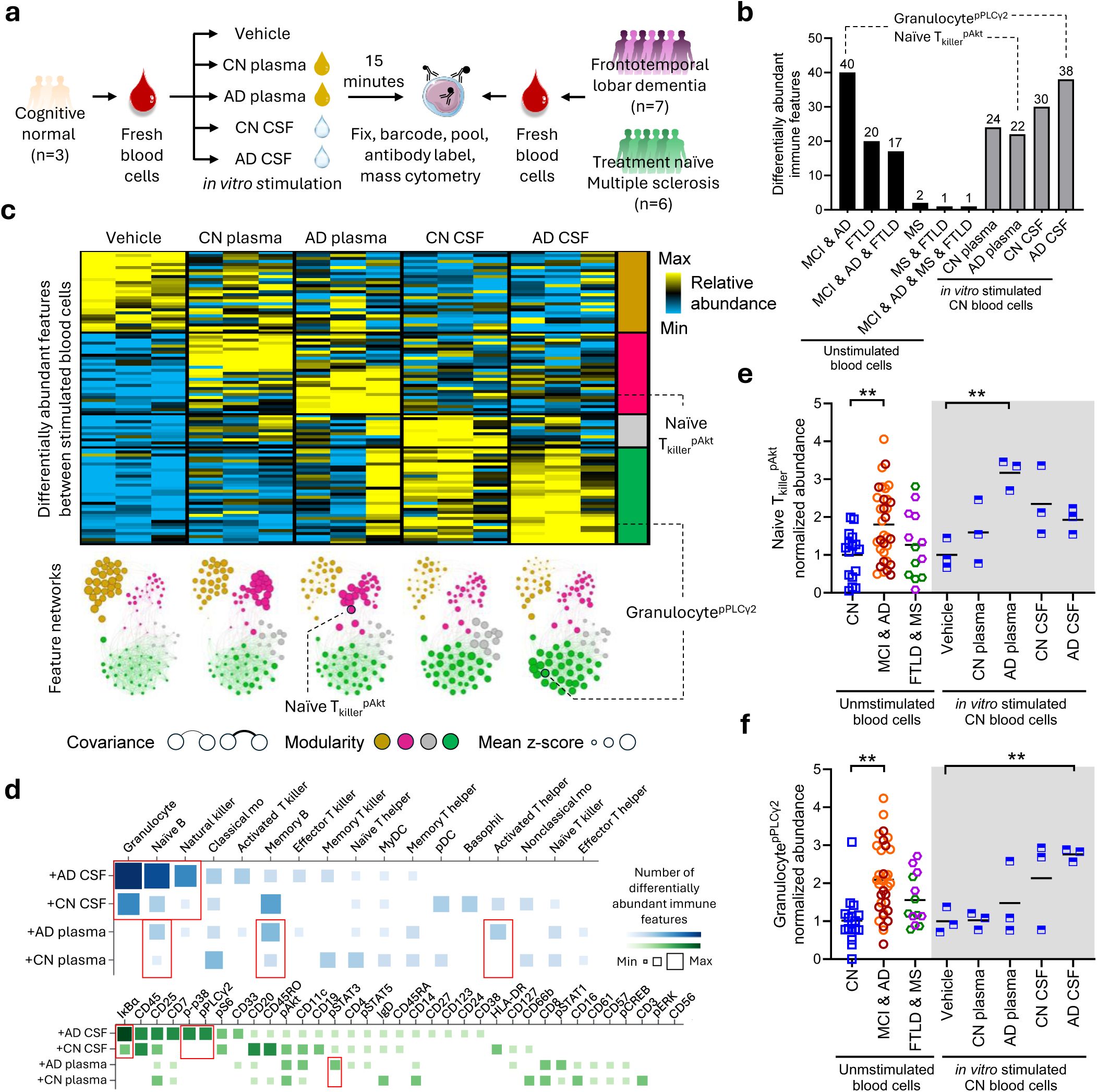
Alzheimer’s disease plasma and cerebrospinal fluid promote disease specific proteomic signatures. **a**, Experimental design. Freshly isolated whole blood cells from three independent cognitively normal donors (biological replicates) were incubated for 15 minutes at 37 °C with plasma or cerebrospinal fluid (CSF) obtained from either cognitively normal or biomarker confirmed diagnosed Alzheimer’s disease donors. Vehicle-treated cells served as additional controls. Following stimulation, cells were immediately fixed and processed for mass cytometry to quantify cell type–specific protein expression and signaling features. In parallel, unstimulated female whole blood from n=7 Frontotemporal lobar degeneration (FTLD) patients and n = 6 treatment-naï ve multiple sclerosis (MS) patients were profiled by mass cytometry. **b**, Differential immune features. (Left) Quantification of unique and shared differentially abundant immune features across female mild cognitive impairment due to Alzheimer’s disease (MCI), Alzheimer’s disease dementia (AD), FTLD and MS groups relative to cognitively normal (CN) controls. (Right) Quantification of unique or shared differentially abundant immune features across indicated *in vitro* stimulated conditions relative to vehicle treated controls. Statistical analysis: two-way ANOVA with Dunnett’s multiple comparison test. Bars represent the number of features with adjusted p-value < 0.05. **c**, (Top) Heat map of z-score–normalized differentially abundant immune features identified in (**b**), averaged across three biological replicates. (Bottom) Network representation of these features. Each node represents a single immune feature and is colored according to modularity class. Node size reflects the mean z-score across replicates. Edge weights correspond to the magnitude of pairwise covariance between features, with edges displayed above a predefined correlation threshold (see Methods). **d**, Cellular and antigen mapping. Matrix plot summarizing differentially abundant immune features stratified by gated immune cell population (top, blue annotation) or by protein marker (bottom, green annotation). Key signaling antigens altered upon exposure to Alzheimer’s disease CSF or plasma are highlighted with red boxes. **e–f**, Selective induction by AD biofluids. (Right, gray shading) Representative immune features selectively induced by Alzheimer’s disease plasma (naï ve T_k_^pAkt^) (**e**) or CSF (granulocyte^pPLCγ2^) (**f**). Statistical analysis: one-way ANOVA with Dunnett’s multiple comparison test versus vehicle control. (Left, unshaded) Normalized signal intensity of corresponding immune features in unstimulated whole blood from female CN (blue), MCI (orange) & AD (maroon), or FTLD (purple) & MS (green) individuals. For all panels, data represent mean from three independent donor replicates unless otherwise indicated. Multiple comparison adjustments were performed as indicated for each panel. *Adjusted p-value < 0.05; adjusted **p-value < 0.01.

To understand whether these immune cell signatures would be present in patients with other neurodegenerative disorders, we performed mass cytometry on unstimulated peripheral blood cells collected from a new cohort of females with either Frontotemporal lobar degeneration (FTLD; n= 7; 62.4±6.2 years of age) or treatment-naï ve multiple sclerosis (MS; n=6; 44.3±15.9 years of age) (**Fig 5a**). We identified immune features that were unique (p-value < 0.05, MCI & AD n=40 features; FTLD n=20 features; MS n=2 features) or shared (MCI & AD & FTLD n=17 features, FTLD & MS n=1 feature, MCI & AD & FTLD & MS n=1 features) across the disease conditions when compared with CN controls (**Fig 5b**). Notably, the AD-related elevation of naï ve T killer^pAkt^ and granulocyte^pPLCγ2^ were not enriched in unstimulated blood cells from patients with FTLD or MS (**Fig 5e-f**). The findings provide direct evidence that soluble factors present in plasma and CSF of AD patients are sufficient to initiate similar peripheral blood immune signaling programs characteristic of AD in naï ve healthy individuals within a short time interval.

## Discussion

In this study, we present a comprehensive analysis of human immune populations at single cell resolution simultaneously obtained from CSF and peripheral blood compartments across the continuum of AD biological and clinical stages and from cognitively normal controls. We note that AD progression triggers peripheral and central immune alterations that follow a predictable, wave-like pattern over time. Increasing AD severity was accompanied by specific and dynamic modifications in innate and adaptive immune system function in both compartments with striking sex differences that were largely independent of *APOE* ε4 genotype status. Our network-based analysis uncovered a set of highly coordinated immune modules whose stage-specific activation patterns correspond to preclinical, MCI and AD dementia stages of disease. Importantly, peripheral immune modules in CN Aβ+ individuals often precede significant changes in plasma ptau_217_, CSF ptau_181_, and BBB permeability, suggesting that systemic immune alterations could potentially be indicators of a biologically distinct early stage of AD progression rather than mere consequences of later neurodegeneration.

Our analyses also revealed that coordinated changes in signaling proteins, rather than immune cell surface receptors, act as central hubs of immune network regulation, reflecting integrated cellular activation across multiple immune cell lineages. Notably, signaling molecules like pPLCγ2, pAkt, pSTAT5, and p-p38 emerged as key drivers of immune coordination, with memory T helper cells consistently exerting strong influence in both central and peripheral immune environments. Alterations in immune signaling pathways may serve as early, system-wide markers of dysregulation in AD, reflecting convergence of multiple upstream signals before changes in immune cell populations or receptor expression become apparent.

Sex was seen as a key determinant of immune homeostasis in both healthy and AD stages, with female immune cells showing higher expression of co-stimulatory and phagocytosis-related markers, and male immune cells enriched for cytotoxic markers. Sex-specific alterations in immune cell state were observed across AD stages, both in peripheral blood and CSF. Taken together, the activation of peripheral signaling pathways are an early event in AD progression and cognitive decline and appear more striking in females than males.

Among females, the robust accumulation of CD33 on monocytes, granulocytes, DCs, basophils and naï ve B cells that we observe in the blood of CN Aβ+ and MCI individuals may represent an early response to amyloidosis. This is consistent with GWAS studies that have identified risk loci within the *CD33* gene locus that are associated with greater cell surface expression of CD33 on monocytes^46^. In contrast, loss of CD33 on the surface of myeloid cells in the AD dementia stage may reflect chronic exposure to oxidative stress, similar to patterns observed in Type 2 diabetes^61^.

Within each sex, immune changes in CSF and blood were concordant, reflecting convergent immune pathways that vary by compartment and disease stage. Notably, peripheral immune alterations were detected prior to the accumulation of ptau_217_, a biomarker with established diagnostic and prognostic value for predicting cognitive decline in cognitively normal, amyloid-positive individuals^62,63^. This temporal relationship highlights the potential utility of peripheral immune proteomic profiling to complement ptau_217_ as a minimally invasive method for early identification of individuals at-risk for AD. Together, these original findings challenge the view of systemic immunity as a passive bystander, instead positioning it as a dynamic and early effector in AD pathophysiology that is modulated across disease stages. Furthermore, these results establish a powerful framework for using immune profiling to enhance diagnostic precision, monitor disease progression longitudinally, and inform development of AD stage specific immunotherapies in the future.

Strengths of this study include a comprehensive single cell proteomic characterization of CSF and peripheral blood immune activation, *APOE* genotype and AD biomarkers, as well as BBB integrity using MRI data and cognitive assessments. Immune alterations were correlated with plasma ptau_217_, CSF Aβ_42/40_ ratios, and cognitive assessments, strengthening their relevance to AD pathophysiology. This systems level analysis allowed for a deeper understanding of immune state changes across the AD clinical spectrum with validation of stronger early activation of the peripheral immune system in females across two clinical cohorts. The results were robust and included integration of multiple computational approaches (SAMuRAI, immune modules, network centrality metrics, PCA, Andrew’s curves, and elastic net regression) to identify features with high confidence. Further, our *ex vivo* stimulation experiments provide mechanistic insight into how systemic immune signatures may arise, further implicating soluble factors in AD plasma and CSF as potentially causal mediators of the immune cell signatures we observe in resting cells from individuals with underlying AD neuropathology.

Limitations of the study include its cross-sectional nature despite profiling individuals in different stages of AD, and the need to further investigate the impact of mixed pathology and multiple medical co-morbidities that are highly prevalent in later stages of AD dementia. An additional limitation is the modest sample size relative to studies that profile proteomic AD biomarkers in biofluids. This can lead to limited interpretation in small subgroups like *APOE* status. However, these limitations were partially mitigated by large effect sizes observed in key findings and by the replication of immune signatures in an independent PBMC CyTOF dataset, which supports the robustness and reproducibility of the results. These findings need future validation in racial and ethnically diverse longitudinal cohorts with functional validation of the immune activation signatures to confirm their exact roles in the ongoing immune response with AD progression.

In summary, this study demonstrates that immune cell activation profiles occur in coordinated waves that are robust in the peripheral blood and CSF in AD biological and clinical stage- and sex-specific-manner. Distinct immune signatures and earlier activation of immune cell signaling pathways in females suggest that the nature of immune response in AD is altered despite similar underlying proteinopathy. Notably, peripheral immune modules in cognitively normal, amyloid-positive individuals emerge prior to detectable changes in plasma ptau_217_, CSF ptau_181_, and blood-brain barrier permeability, suggesting that systemic immune alterations may reflect a biologically distinct early stage of disease progression rather than a subsequent byproduct of neural degeneration. Together, these findings position peripheral immunity as a dynamic, stage-specific actor in AD pathophysiology and highlight candidate peripheral and CNS immune signatures for mechanistic studies and their potential for immune profiling for improving early detection, disease monitoring and guiding future immunotherapeutic strategies.

## Author contributions

Conceptualization: A.B., M.L., H.M., W.B. and J.A.P

Clinical evaluation and coordination: Q.B., St.R., D.O., J.B.L., and J.A.P.

Sample collection and fluid biomarker analysis: L.M.B., Q.B, Sa.R., E.W., and M.K.

Mass Cytometry: H.M.

DCE-MRI: M.L. and W.S.

Data Visualization and statistical analysis: A.B., B.M., S.C., Sa.R., W.B., P.B., J.A.P.

Funding acquisition: J.A.P, A.B., W.B., M.L., W.S., H.M., St.R., D.O., J.B.L.

Writing – original draft: A.B. and J.A.P.

Writing – review & editing: J.A.P, A.B., M.L., W.S., B.M., H.M., L.M.B, Q.B., S.C., Sa.R., E.W., M.K., W.B., P.B., St.R., D.O., J.B.L.

## Data and materials availability

All data associated with this study are present in the paper or the Supplementary materials. Newly generated mass cytometry data files will be made available through Immport SDY3416.

## Materials and Methods

### Participants

The study was approved by the Cleveland Clinic Institutional Review Board. All participants provided written, informed consent. The prospective cohort included 70 individuals (> 60 years old) evaluated at the Cleveland Clinic Center for Brain Health between 2023 and 2024. All participants were fluent in English and included HC, MCI from underlying AD, and AD dementia individuals. Individuals considered CN had a Mini Mental State Examination (MMSE) score of >25, a clinical dementia rating-global (CDR-G)^42^ score of 0, no subjective memory concerns, biomarkers (Aβ42, total tau (t-tau), phospho tau181 (ptau_181)_) in their CSF not supportive of underlying AD biological changes and were recruited from the general community. MCI and AD dementia individuals were recruited from a specialized memory clinic at Cleveland Clinic and were evaluated by a specialized physician with expertise in dementia care. MCI individuals met NIA/AA criteria for MCI from AD^64,65^ and had a CDR-G score of 0.5. AD dementia individuals met NIA/AA criteria for dementia from AD^65,66^ and had a CDR-G score of ≥1. The presence of AD biomarkers among the MCI and AD dementia individuals were confirmed after CSF evaluation of Aβ42,40, t-tau and ptau_181_ (Table 1). Participants also completed the Montreal Cognitive Assessment (MoCA)^67^ and Clinical dementia rating scale sum of boxes (CDR-SB)^42^ for cognitive and functional evaluation.

Participants in the Frontotemporal lobar degeneration cohort were diagnosed by a clinical consensus panel diagnosis after review by neurologists/neuropsychologist per current FTLD diagnostic criteria^68–71^ and negative for plasma AD biomarkers (Aβ42/40 ratio and ptau 217). FTLD participants also completed the Montreal Cognitive Assessment^67^.

Participants in the multiple sclerosis cohort were diagnosed according to established clinical criteria^72^ from a multidisciplinary referral MS center at the Cleveland Clinic. Individuals were eligible for inclusion if they were 25 years or older and had a confirmed diagnosis of MS documented in their medical records. To minimize potential confounding effects of pharmacologic treatments on study outcomes, only individuals who were treatment naï ve for disease-modifying therapies (DMTs) were included. In addition, participants were required to have no exposure to systemic corticosteroids within the previous six months prior to enrollment. MS participants also completed the Montreal Cognitive Assessment^67^.

### Additional inclusion/exclusion criteria

Participants were excluded if they have a history or evidence of:

1) neurological illnesses/conditions (other than MCI, FTLD or MS for those specific cohorts), such as Parkinson’s, or Huntington’s disease, head trauma with significant loss of consciousness (>30 min) or residual neurological deficit, cerebral ischemia, vascular headache, carotid artery disease, cerebral palsy, epilepsy, brain tumor, chronic meningitis, multiple sclerosis, pernicious anemia, normal-pressure hydrocephalus. 2) medical illnesses/conditions that may affect brain function, such as untreated hypertension (blood pressure >140/100 mm Hg), unstable cardiac disease, uncontrolled treated insulin-dependent diabetes mellitus and other endocrine disorders, severe renal disease, severe glaucoma, and chronic obstructive pulmonary disease. 3) current Axis I psychiatric disturbance meeting DSM-IV Axis I criteria. 4) severe depressive symptoms. 5) substance abuse or dependence. 6) exclusion criteria specific to MR scanning (pregnancy, weight inappropriate for height, ferrous objects within the body, low visual acuity, and a history of claustrophobia); 7) current use of psychoactive medications, except SSRI and SNRI antidepressants. 8) any unstable or severe cardiovascular disease or asthmatic condition. 9) active inflammatory disorder or infections after screening lab tests. 10) history of imaging confirmed transient ischemic attack or a score of >4 on the modified Hachinski ischemic scale. 11) history of coagulopathy or use of anticoagulants contraindicated for lumbar puncture. 12) exposure to daily nonsteroidal anti-inflammatory (NSAID), steroids, biologics, chemotherapy medications, but not daily Aspirin (81 mg). 13) hospitalization in the last 6 months for surgeries and infections.

### Biosamples for AD biomarkers and APOE genotype

All blood and CSF samples are collected in the morning after overnight fasting. Samples were frozen within 15 min of collection at −70°C and evaluated after one freeze-thaw cycle at the time of analysis. Assays used comparable lumbar puncture fractions to limit variability from rostrocaudal concentration gradients. CSF levels of Aβ42, t-tau, and ptau_181_ levels, were measured using MILLIPLEX MAP® multiplex kits and analyzed using a Luminex 200 3.1 xPONENT System (Luminex xMAP technology; EMD Millipore, Chicago). All samples were analyzed blind to diagnosis. Additionally, plasma Aβ40, Aβ42, and ptau_181_ were analyzed using a Simoa Assay (Quanterix, Billerica, MA). Assays for plasma ptau_217_, was completed using a SinglePlex Assay (MSD, Rockville, MD). Apolipoprotein E (*APOE*) genotyping from blood samples was performed using the 7500 Real Time PCR System and TaqMan SNP Genotyping Assays (rs429358, rs7412) (Thermo Fisher Scientific) as previously described^73^.

### MRI Scanning

All scanning was performed on a Siemens 3T MRI scanner with a 32 receive channel head coil. The following scans were obtained: A) Anatomical Imaging using a 3D T1-Weighted Scan (MPRAGE, FOV = 256 x 256 mm2, 120 sagittal slices, resolution = 1 x 1 x 1.2 mm3, TI/TR/TE/flip angle = 900/1900/1.7 ms/8°), a 2D T2-Weighted Scan (TSE, resolution = 0.7 x 0.7 x 4 mm^3^, TR/TE1/TE2 = 2000/20/110 ms, FOV = 210 x 210 mm^2^, 35-4 mm axial slices). Hippocampal and cortical volumes were determined using Freesurfer v7.1.0^74^. Dynamic contrast-enhanced (DCE) MRI using 3D T1w gradient refocused echo (GRE) acquisition [flip angle = 15°, voxel size = 1.9x1.9x5 mm^3^, 16 slices, 6.2 s of temporal resolution (time per volume acquisition)] over 14 minutes was obtained using a 0.1 mmol/kg dosage of Gadolinium contrast (Dotarem, Buerbet, France, r1=4.5 s^-1^mM^-1^ at 3T) injected through intravenous bolus injection at 4 ml/s for 20 s after the 5^th^ volumes of DCE scan followed by an injection of 20 ml of saline at the same rate. The Patlak model^75^ has been shown to produce reliable results in prior studies of neurodegeneration^76,77^ and was used to calculate permeability. Prior to DCE MRI, two additional spoiled GRE scans with different flip angles (5 and 8 degrees) were collected for T1 calculation^78^. Analysis was conducted using open source DCE analysis toolbox^79^. Permeability in anatomically segmented regions using public open software Freesurfer v7.1.0^74^ was assessed including the hippocampus.

### Whole blood stimulation with plasma or CSF prior to mass cytometry evaluation

Human whole blood from cognitively normal individuals was collected into sodium heparin coated tubes (Greiner Vacuette) and stored on wet ice for up to two hours. 400 µL of blood per condition was passed through a 40 micron filter into a 5 mL Falcon round bottom tube (Corning) and allowed to rest at 37°C 5% CO2 for 25 minutes. Flash-frozen human plasma or CSF from cognitively normal or Alzheimer’s disease dementia individuals was thawed in a 37°C bead bath to be used for stimulations. Rested blood tubes were brought into a biosafety hood and 100 µL of sterile PBS, 100 µL plasma or 100 µL CSF added to appropriate tubes and pipetted up and down three times to mix. Samples were immediately placed in 37°C incubator to allow for signaling to occur. After exactly 15 minutes, 700 µL of PROT1 Proteomic Stabilizer (Smart Tube Inc, San Carlos, CA) was added to each stimulated blood sample, followed by pipetting up and down four times. Cells were allowed to fix at room temperature for 10 minutes, then transferred to cryovials and frozen at −80°C. Samples were processed by mass cytometry as described below for unstimulated cells.

### Phosphoflow whole blood and CSF mass cytometry

CSF without blood contamination was collected by lumbar puncture following local anesthesia and was centrifuged at 400xg for 10 minutes at room temperature within 1-2 hours of collection. Following reservation of CSF supernatant for soluble analyte assessment, cell pellets were resuspended in 500 µL residual volume and added to a barcoded tube containing 700 µL of PROT1 Proteomic Stabilizer (Smart Tube Inc, San Carlos, CA). After 10 minutes at room temperature, the tube was placed on dry ice and stored at −80°C. Whole blood was collected into EDTA coated tubes and 500 µL of blood was fixed in 700 µL of PROT1 Proteomic Stabilizer for 10 minutes at room temperature, then the tube was placed on dry ice and stored at −80°C. Cell staining and mass cytometry was performed by the Human Immune Monitoring Center at Stanford University. Upon thawing, 200 µL aliquots of whole blood or CSF were plated per well in 96-well deep-well plates or cluster tubes. After resting for 15 minutes at 37°C, samples were washed with Smart Tube Thaw-Lyse buffer (Smart Tube Inc.) twice, and with CyFACS buffer (PBS supplemented with 2% BSA, 2 mM EDTA, and 0.1% sodium azide) once, then permeabilized with 100% ice cold methanol and kept at −80°C overnight. The next day cells from up to 20 samples, balanced for age, sex, and disease status, were barcoded using Cell-ID™ 20-Plex Pd Barcoding (Standard Biotools, Markham, Canada), followed by staining for 30 min at room temperature with 20 mL of surface and intracellular antibody cocktail. Cells are then washed twice with CyFACS and resuspended in 100 mL iridium-containing DNA intercalator (1:2000 dilution in 2% PFA in PBS) and incubated at room temperature for 20 min. Cells are then washed once with CyFACS buffer and twice with MilliQ water, then diluted to 7.5x10^5^ cells/mL in MilliQ water and acquired on Helios CyTOF. Quality control was performed using FlowJo v10 by gating on intact cells based on the iridium isotopes from the intercalator, then on singlets by Ir191 vs cell length followed by cell subset-specific gating.

### Elastic net

For Plasma p-tau217, we fit elastic net regression models using the R v4.5.1 package glmnet v4.1-10 assuming a Gaussian outcome. Mixing parameters α were evaluated over a grid from 0 to 1 in increments of 0.05. For each fixed α, we used 5-fold cross-validation (cv.glmnet) to select the regularization parameter λ as the value that minimized the mean cross-validated mean squared error (MSE). The final model was selected as the (α, λ) combination achieving the lowest cross-validated MSE across the α grid. For Status, as defined in the Recruitment section, we fit elastic net–regularized cumulative logit (proportional-odds) models using the R v4.5.1 package ordinalNet v2.13 with standardized predictors and evaluating a grid of mixing parameters, α, ranging from 0 to 1 in increments of 0.25. For each α, we used 3-fold cross-validation (ordinalNetTune) to select the regularization parameter λ by maximizing the cross-validated mean log-likelihood. We selected the final (α, λ) as the combination with the highest cross-validated mean log-likelihood.

### Statistical Analyses

Most of the data were analyzed using Graphpad Prism 10 (GraphPad Inc). For comparisons across multiple conditions, one- or two-way ANOVA with Sidak, Tukey’s or Dunnett’s correction for multiple comparisons. For distributions that did not meet the assumption of normality using the Shapiro-Wilk test, we applied Kruskal-Wallis test with Dunn’s multiple comparisons test. We calculated mean values for continuous variables between the three groups using a two-way Analysis of variance (ANOVA) with Dunnett multiple comparison test p < 0.05. Tukey multiple comparison testing was used when evaluating multiplex analytes and immune feature abundance. Unsupervised hierarchical clustering was performed using the One minus Pearson correlation tool (https://software.broadinstitute.org/morpheus). Network plots were generated using Gephi.

---

**ED Fig. 1.**
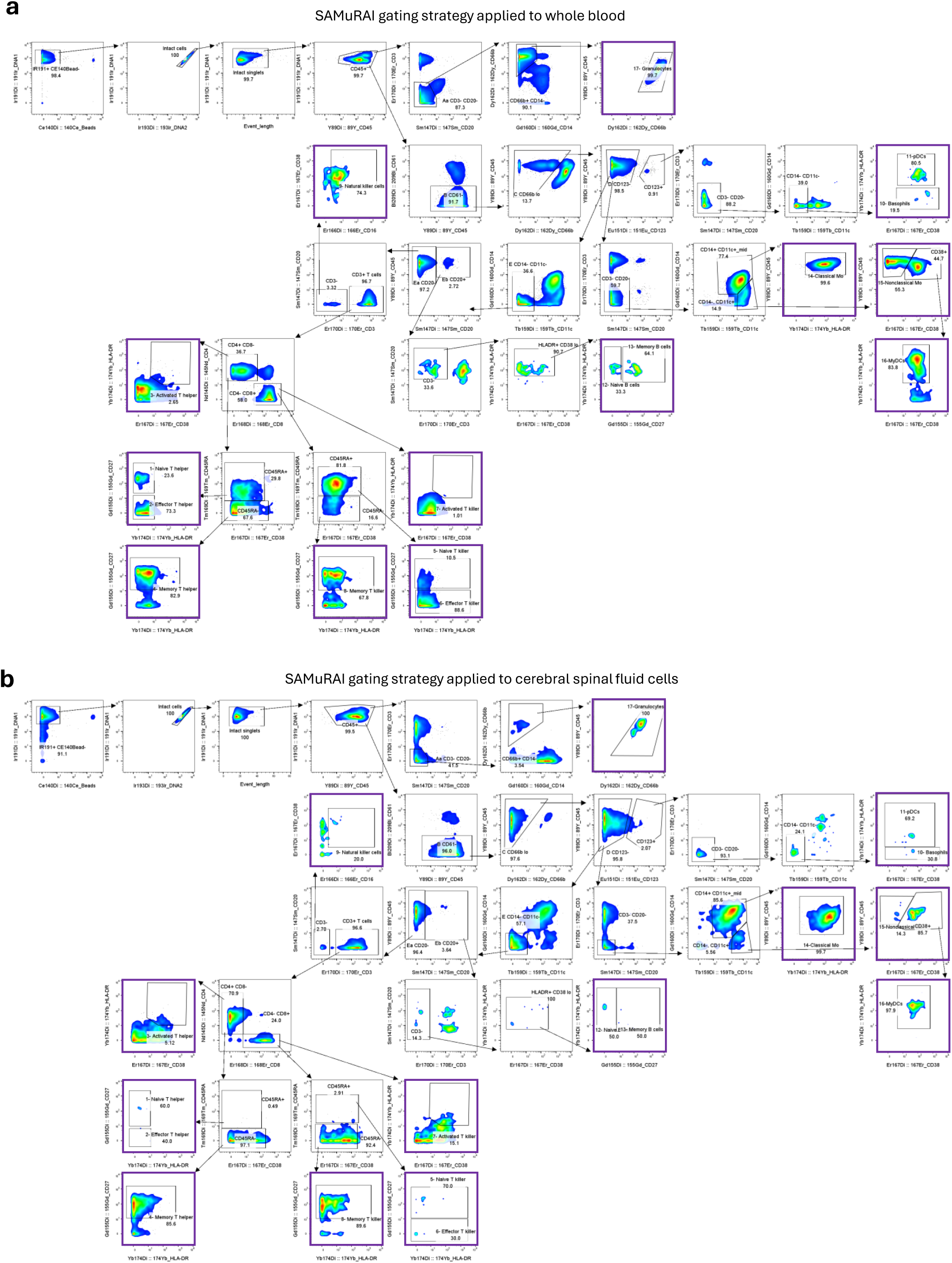
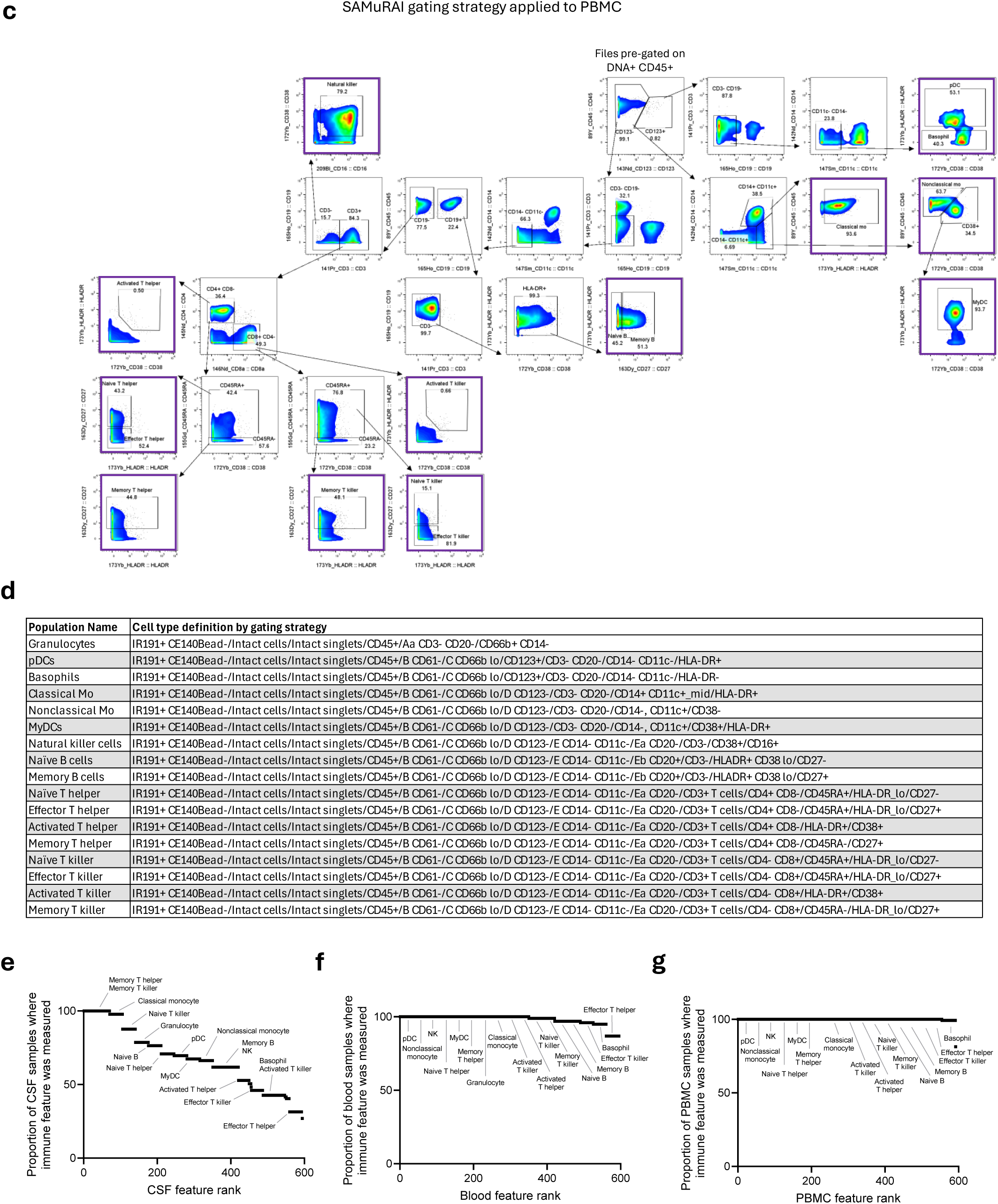
Gating scheme to identify major cell populations in peripheral blood, CSF, and PBMC. **a-c**. A common gating scheme was applied to peripheral blood (**a**), CSF (**b**) or PBMC (**c**) mass cytometry files to identify 17 major immune cell types, except granulocytes which are depleted during the PBMC isolation procedure. **d,** Definitions of cell populations by the combination of surface antigens used. **e-f**, Proportion of (**e**) CSF (**f**) blood or (**g**) PBMC samples in which immune features were determined. When a gated cell population was not detected in a sample, the immune features from that population could not be determined.

**ED Fig. 2.**
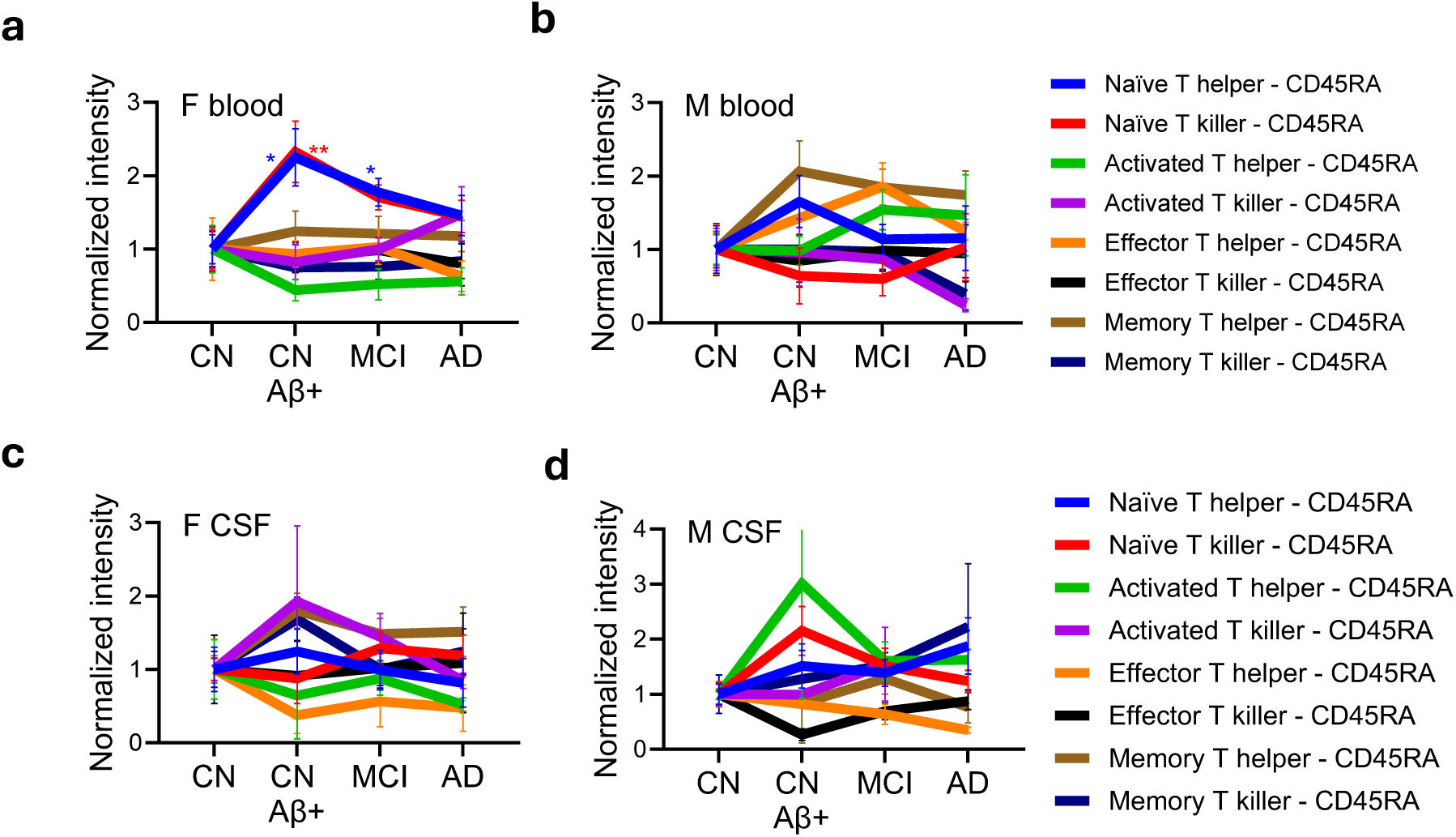
CD45RA relative intensity in blood and CSF T cell population across AD stages. **a-b**, Intensity of CD45RA in circulating T cells subsets in females (**a**) or males (**b**). **c-d**, Intensity of CD45RA in CSF T cells subsets in females (**c**) or males (**d**). * p < 0.05 ** p < 0.01. Two-way ANOVA with Dunnett multiple comparison.

**ED Fig. 3.**
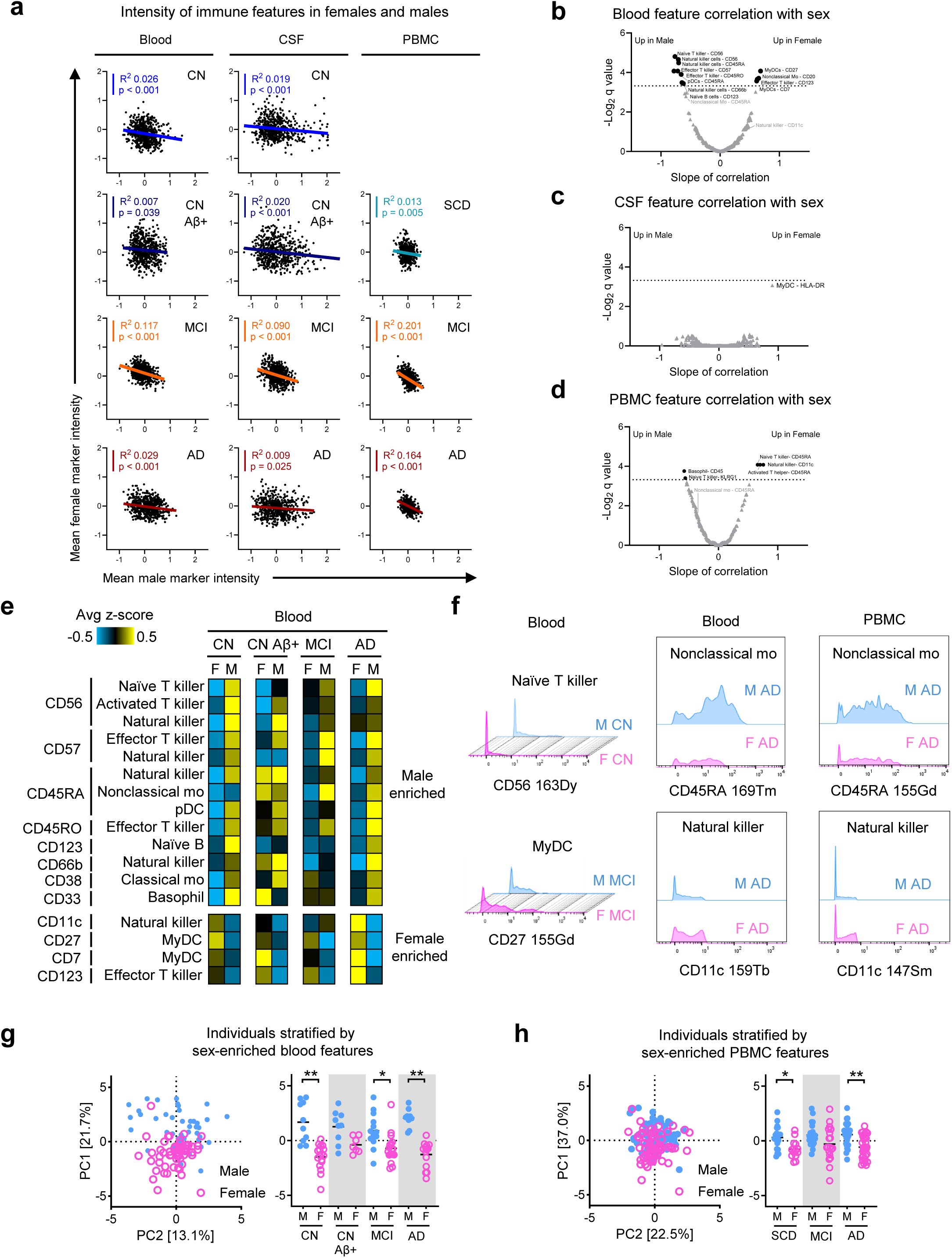
Mass cytometry identifies sex-enriched peripheral immune cell features. **a,** Average z-score normalized intensity of immune features in females (y-axis) or males (x-axis) derived from blood, CSF, or PBMC fractions stratified by AD stage. Each dot represents 595 immune features in blood and CSF or 592 features in PBMC. Bold lines represent linear regression of all markers (colored lines by disease stage blue-CN, dark blue-CN Aβ+, orange-MCI, maroon-AD). **b-d**, Volcano plot of Benjamini-Hochberg corrected q value (y-axis) and slope of linear regression (x-axis) of immune features from blood (**b**), CSF (**c**) or PBMC (**d**) relative to sex of donor. **e,** z-score normalized intensity of sex-enriched immune features from blood. **f,** Representative histogram of male-enriched (naï ve T killer^CD56^; nonclassical mo^CD45RA^) or female-enriched (myDC^CD27^) features in blood or PBMC.**g**, Principal component analysis of male (blue) and female (pink) blood donors using features in (**b**). Each dot represents one blood donor sample. Kruskal-Wallis test with Dunn’s multiple comparison test of values from Principal component 1 (PC1). * p < 0.05 ** p<0.01. **h**, Principal component analysis of male (blue) and female (pink) PBMC donors using nonclassical mo^CD45RA^, NK^CD11c^, classical mo^CD38^ and basophil^CD33^. Each dot represents one PBMC donor sample. One-Way ANOVA with Sidak multiple comparison test of values from Principal component 1 (PC1). * p < 0.05 ** p<0.01.

**ED Fig. 4.**
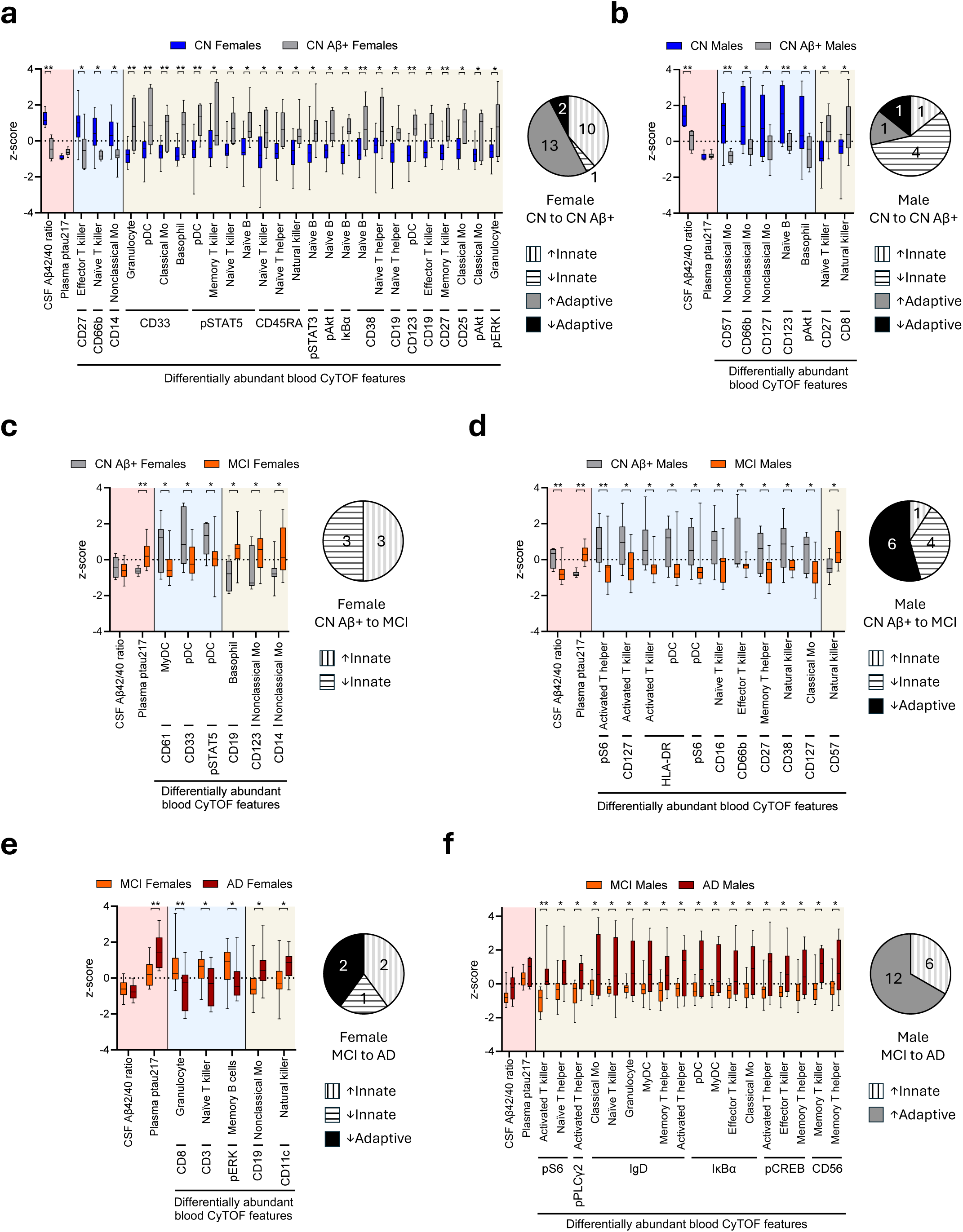
Peripheral immune features dysregulated between AD biological and clinical stages. Related to Fig 1. **a-f**, Blood CyTOF derived immune features that were found to be differentially abundant (adjusted p-value < 0.05) by Two-way ANOVA with Tukey multiple comparison in the analysis performed in **Fig 1c**. Boxes represent 5-95% confidence intervals and whiskers represent max and min values per feature. z-scores of CSF Aβ_42/40_ ratio and plasma ptau_217_ are plotted with light red shaded background in plots for comparison with z-scores of downregulated blood features (light blue shaded background) and upregulated blood features (light tan shaded background). Pie charts represent the number of differentially abundant features within each comparison that included elevated innate immune features (vertical bar white background), reduced innate immune features (horizontal bar white background), elevated adaptive immune features (dark grey background) and reduced adaptive immune features (black background). **a, b,** Differentially abundant blood features between CN (dark blue) and CN Aβ+ (grey) females (**a**) or males (**b**). **c, d,** Differentially abundant blood features between CN Aβ+ (grey) and MCI (orange) females (**c**) or males (**d**). **e, f,** Differentially abundant blood features between MCI (orange) and AD dementia female (**e**) or males (**f**).

**ED Fig. 5.**
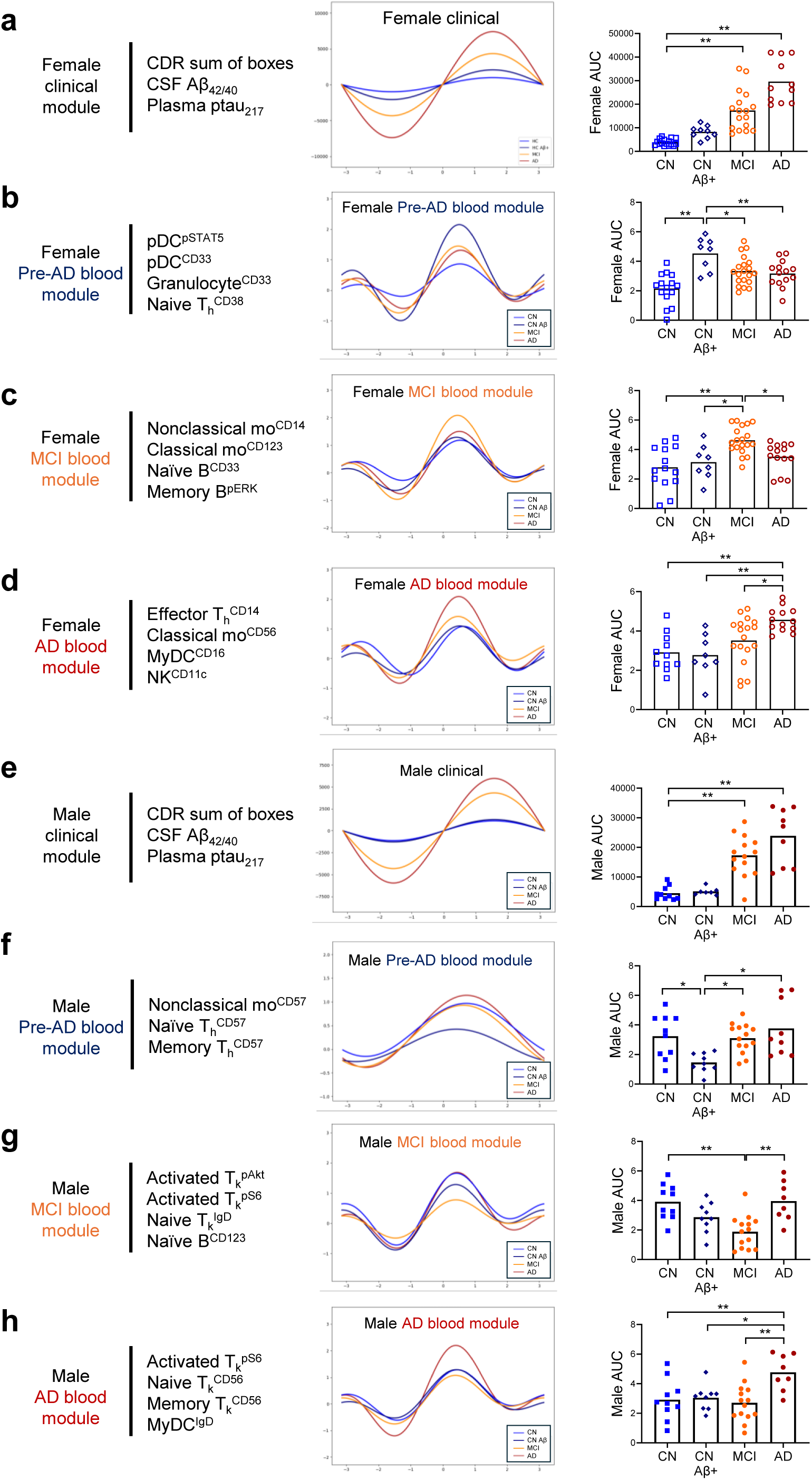
Andrews curves of peripheral blood immune cell states stratify individuals along the AD continuum. Related to Fig 1. **a-h**, Andrews curves were fit for females (**a-d**) or males (**e-h**) using clinical characteristics (**a, e**) or blood immune features (**b-d**, **f-h**). Plots represent the average Andrews curve for each AD stage (CN blue, CN Aβ+ dark blue, MCI orange, AD maroon). Area under the Andrews curve AUC) was calculated for each individual and represented by colored dots in the bar plots. **a-b** and **d-h** One-way ANOVA with Dunnett multiple comparison. **c**, Kruskal-Wallis with Dunn’s multiple comparison. * p < 0.05 ** p < 0.01.

**ED Fig. 6.**
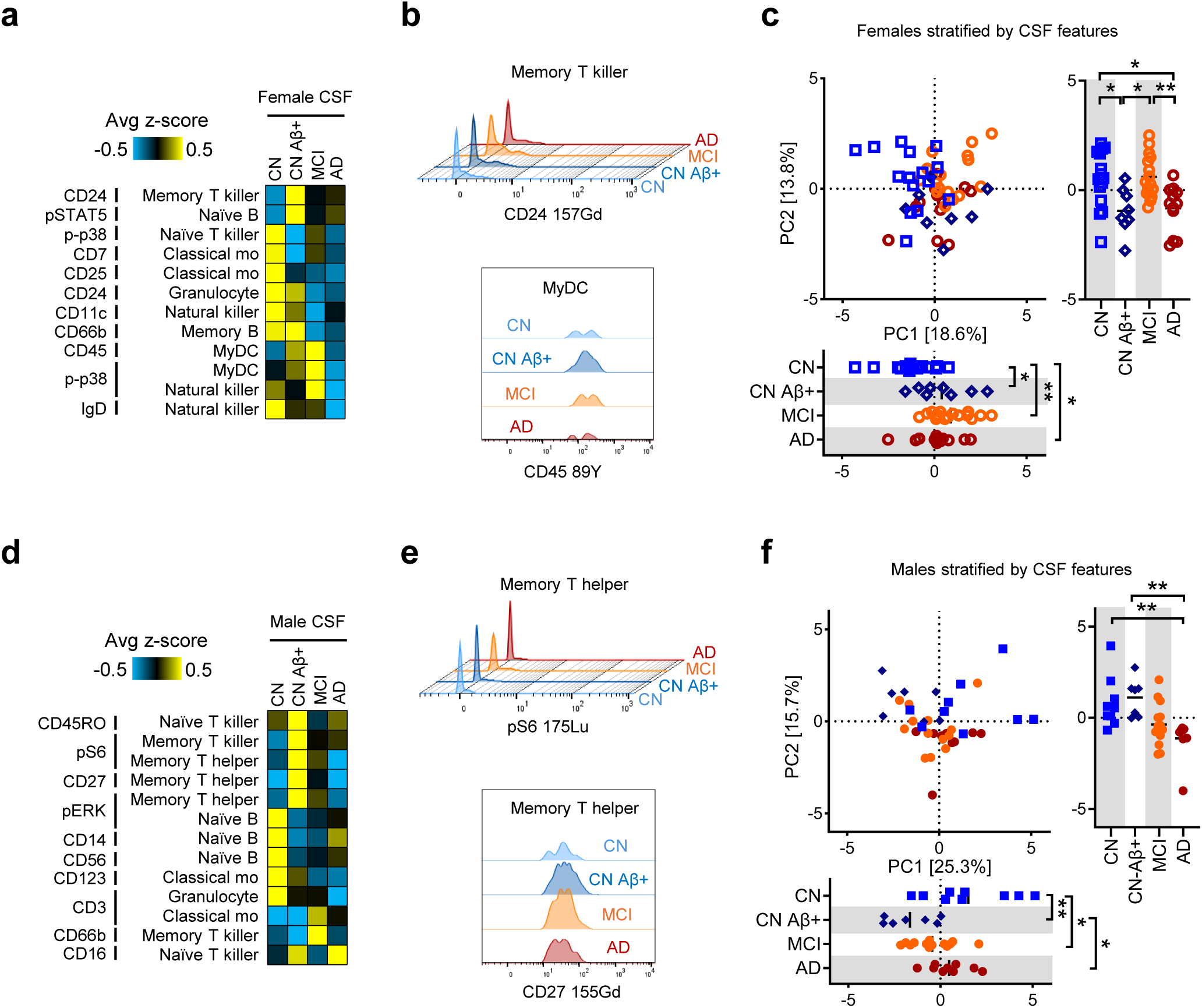
CSF cell states stratify individuals along the AD continuum. Related to Fig 1. **a**, Heat map of average z-scores for female CSF markers that exhibited dynamic expression along the AD continuum. **b**, Representative histogram of female CSF markers that were enriched in CN Aβ+ (Memory T killer^CD24^) or both CN Aβ+ and MCI (myDC^CD45^) stages. **c**, PCA of females using markers in (**a**). Each dot represents one CSF donor sample. One-Way ANOVA with Tukey multiple comparison test of values from PC1 or PC2. * p<0.05 ** p<0.01. **d**, Heat map of average z-scores for male CSF markers that exhibited dynamic expression along the AD continuum. **e**, Representative histogram of male CSF markers that were enriched in CN Aβ+ stage (memory T helper^pS6^; memory T helper^CD27^). **f**, PCA of males using markers in (**d**). Each dot represents one blood donor sample. One-Way ANOVA with Tukey multiple comparison test of values from PC1. Kruskal-Wallis test with Dunn’s multiple comparison test of values from PC2. * p<0.05 ** p<0.01.

**ED Fig. 7.**
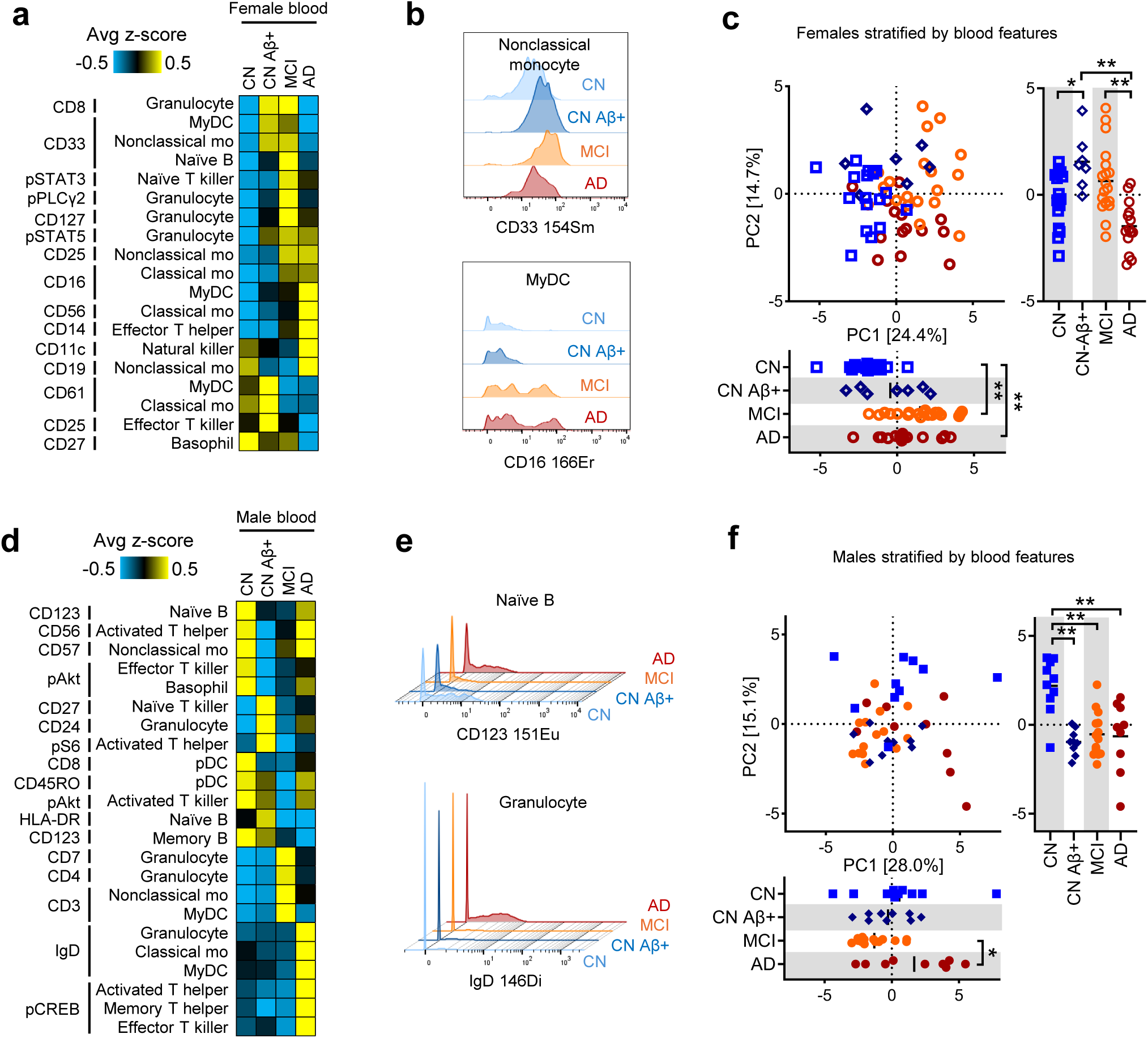
Peripheral blood immune cell states are sufficient to stratify individuals along the AD continuum. Related to Fig 1. **a**, Heat map of average z-scores for female peripheral blood markers that showed dynamic expression along the AD continuum. **b**, Histogram of representative female blood markers that were enriched in CN Aβ+ (nonclassical monocyte^CD33^) or enriched MCI and AD (myDC^CD16^). **c**, PCA of females using markers in (**a**). Each dot represents one blood donor sample. One-Way ANOVA with Tukey multiple comparison test of values from PC1 or PC2. * p<0.05 ** p<0.01. **d**, Heat map of average z-scores for male peripheral blood markers that showed dynamic expression along the AD continuum. **e**. Histogram of representative male blood markers that were depleted in CN Aβ+ and MCI (naï ve B^CD123^) or enriched in AD (granulocyte^IgD^). **f**, PCA of males using markers in (**d**). Each dot represents one blood donor sample. One-Way ANOVA with Tukey multiple comparison test of values from PC1 or PC2. * p<0.05 ** p<0.01.

**ED Fig. 8.**
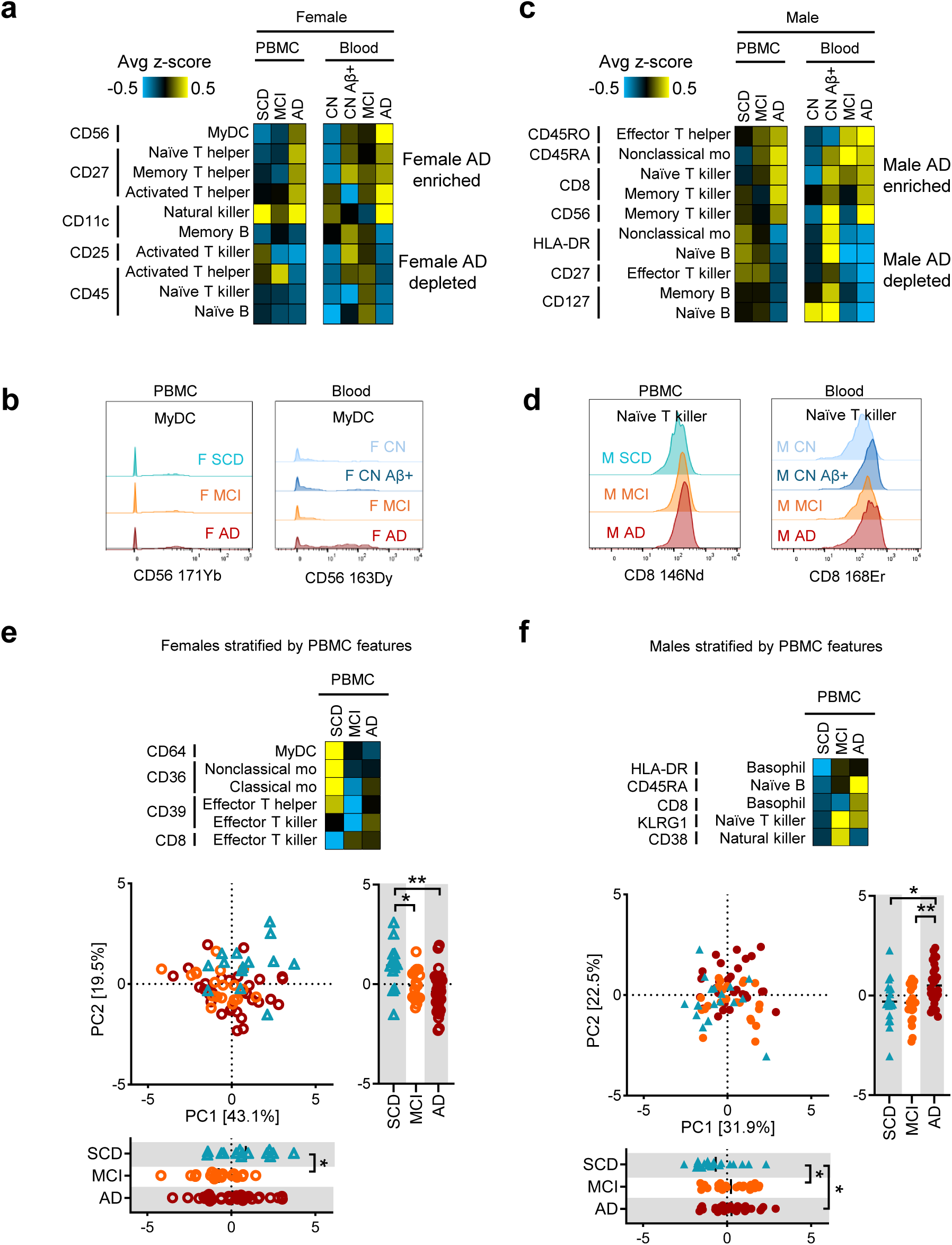
Concordant peripheral immune changes in AD blood and PBMC. Related to Fig 1. **a,** Heat map of average z-scores for female PBMC and blood markers that were concordantly elevated or reduced in AD. **b**, Representative histogram of myDC^CD56^ enrichment in PBMC and blood from AD females. **c,** Heat map of average z-scores for male PBMC and blood markers that were concordantly elevated or reduced in AD. **d**, Representative histogram of naï ve T killer^CD8^ enrichment in PBMC and blood from AD males. **e-f,** Heat map of female PBMC features (**e**) or male PBMC features (**f**) used for PCA. Each dot is a PBMC donor. One-way ANOVA with Tukey multiple comparison. *p<0.05 **p<0.01.

**ED Fig. 9.**
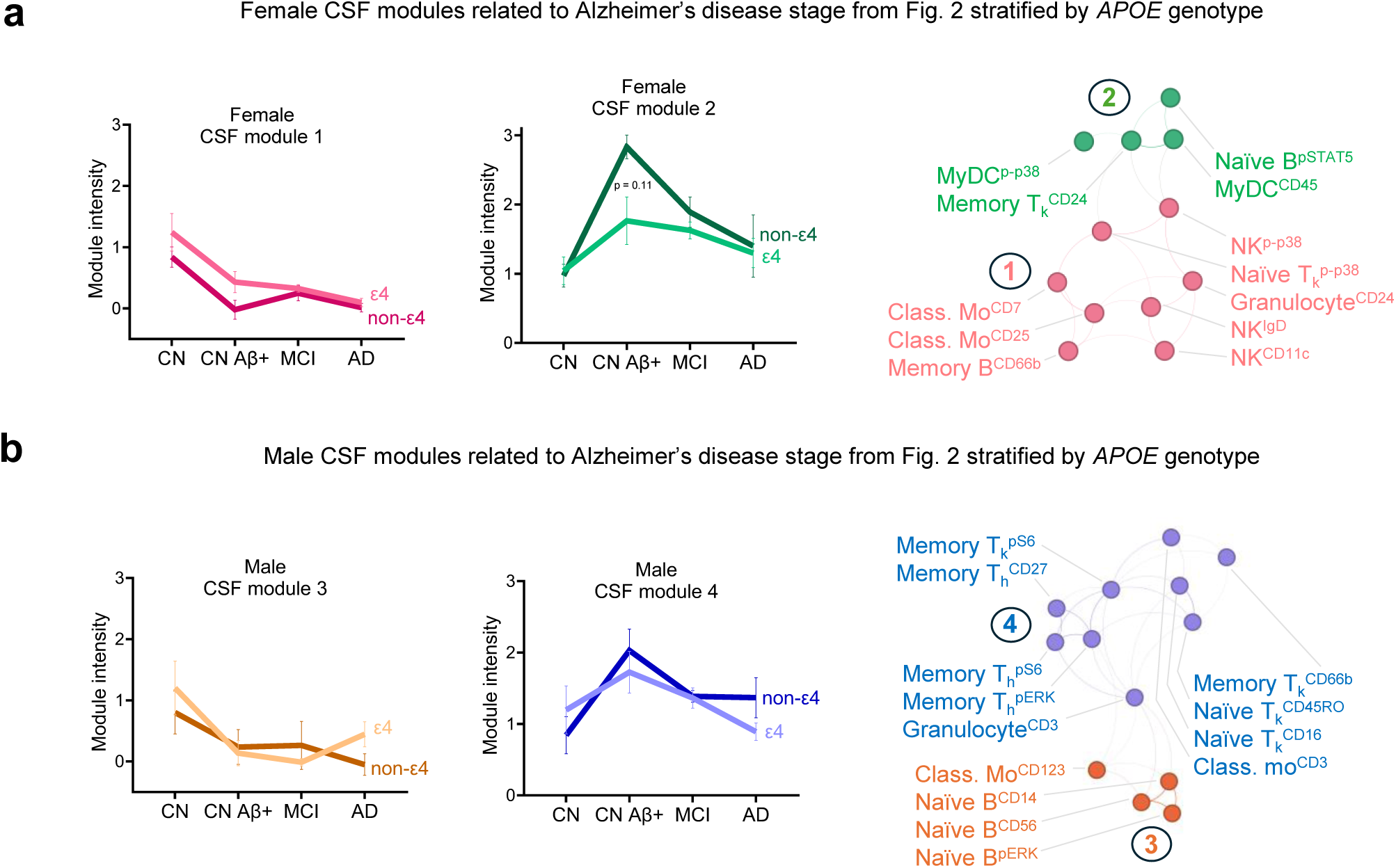
Dynamic central nervous system immune modules stratified by *APOE* genotype. Related to Fig 2. **a-b,** CSF immune modules from females (**a**) or males (**b**) stratified by *APOE ε4* carriers (*ε4*) and non *ε4* carriers (non-*ε4*). Error bars represent S.E.M. No differences reached statistical significance by Two-way ANOVA with Sidak’s multiple comparison test within each disease stage.

**ED Fig. 10.**
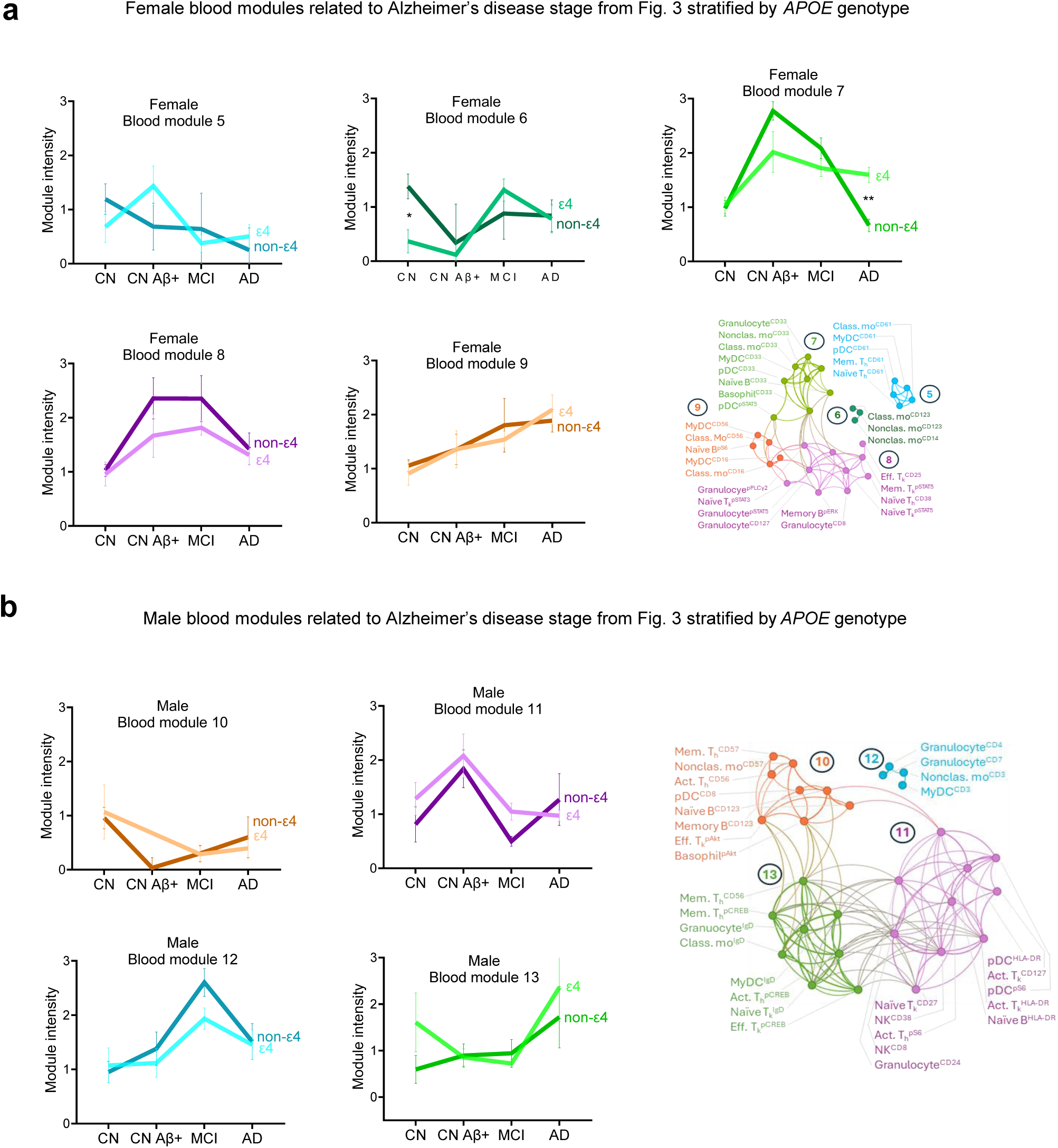
Dynamic peripheral immune modules stratified by *APOE* ε4 genotype. Related to Fig 3. **a-b,** Peripheral blood immune modules from females (**a**) or males (**b**) stratified by *APOE ε4* carriers (*ε4*) and non *ε4* carriers (non-*ε4*). Error bars represent S.E.M. Two-way ANOVA with Sidak’s multiple comparison test within each disease stage. * p < 0.05 ** p < 0.01.

**ED Fig. 11.**
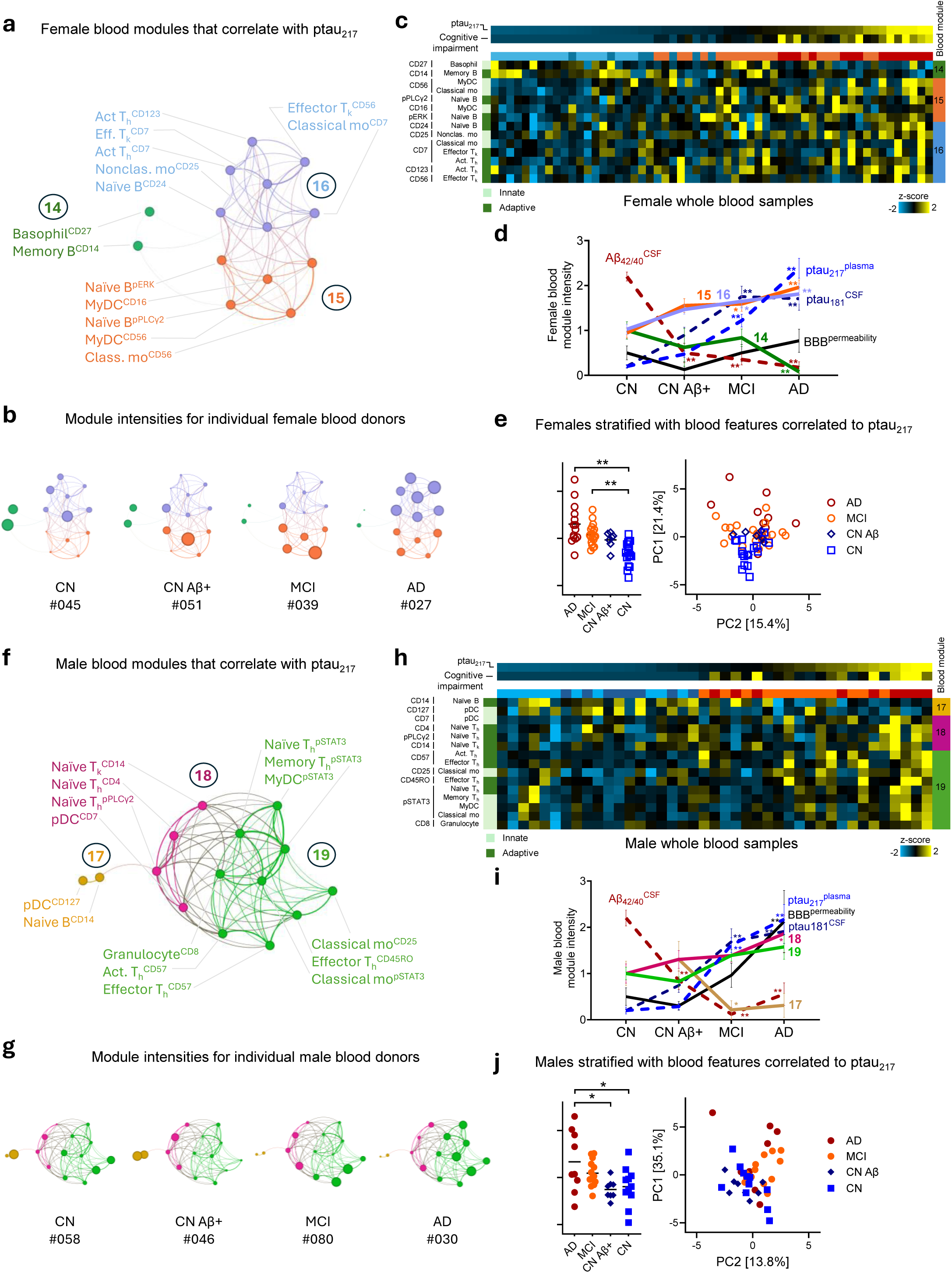
Peripheral immune networks correlate with plasma ptau_217_. **a-i,** Network of female (**a-d**) or male (**f-i**) peripheral blood immune features that correlated with plasma ptau_217_ levels, colored by module with connections representing the strength of feature covariance. **b, g**, Network node sizes were scaled by marker intensity for representative individuals classified by disease stage and patient identifier number. **c**, **h**, Intensity of module components across female (**c**) or male (**h**) blood donors colored by z-score normalized intensity (teal low; yellow high). Plasma phophso-tau_217_ and cognitive impairment represented by Clinical dementia rating sum of boxes are reported above. **d, i**, Summary of module dynamics across disease stages. Hippocampal blood brain barrier (BBB) permeability gleaned from DCE-MRI. Error bars represent S.E.M. Two-way ANOVA with Dunnett’s multiple comparison test relative to CN. * p < 0.05 ** p < 0.01. **e, j** PCA of females (**e**) or males (**j**) using markers in (**a**) or (**f**), respectively. Each dot represents one blood donor sample. One-Way ANOVA with Tukey multiple comparison test of values from PC1. * p<0.05 ** p<0.01.

**ED Fig. 12.**
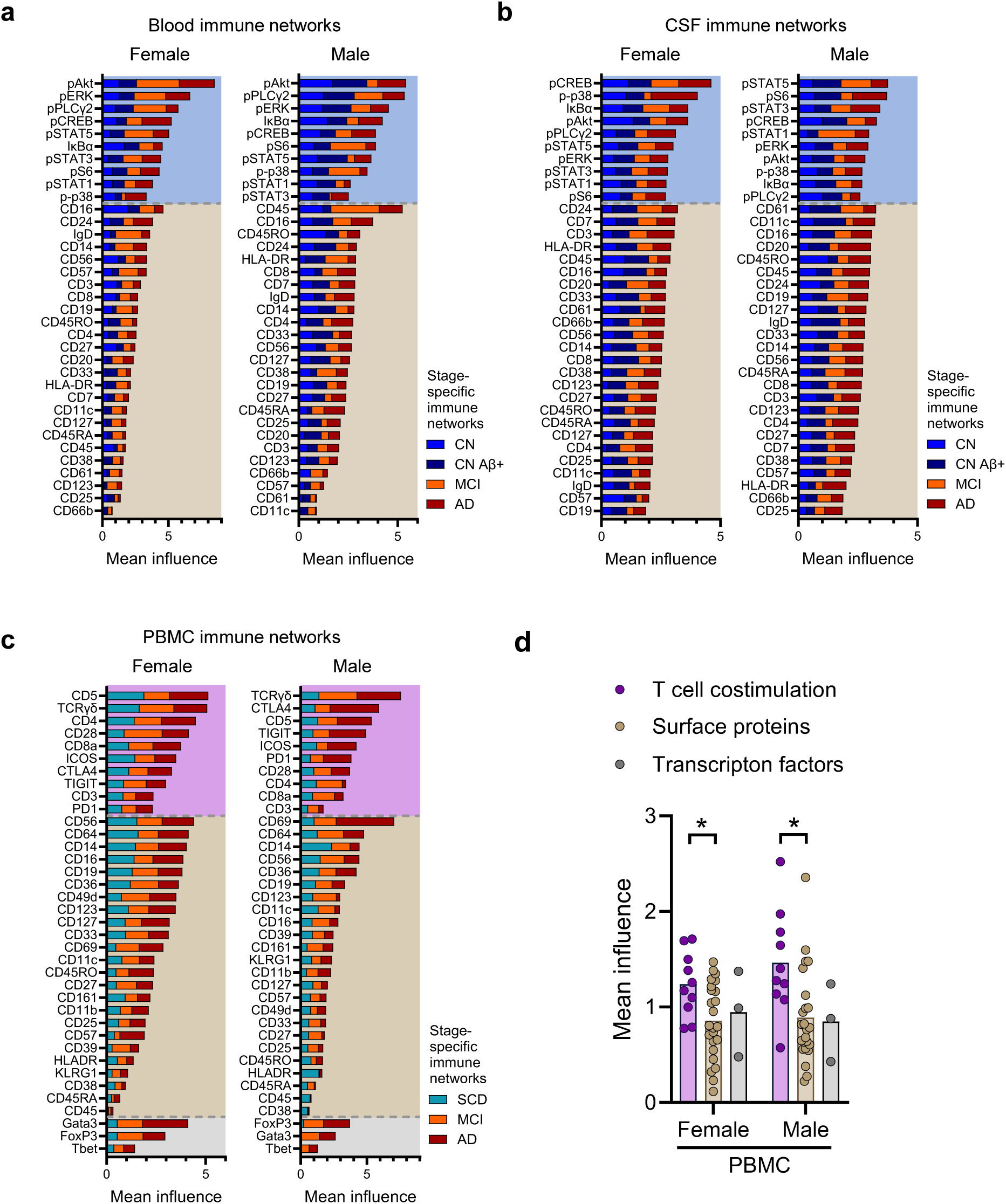
Immune feature influence across clinical stages of Alzheimer’s disease. Related to Fig 4. **a-c,** Feature influence within each sex-stratified tissue-specific network broken out by disease stage-specific networks derived from the covariance networks in Fig 4a-b. Female and male covariance network influence derived from (**a**) blood, (**b**) CSF, or (**c**) PBMC datasets. **d**, Mean influence of PBMC immune features classified T cell costimulation (purple), surface proteins (brown) or transcription factors (grey). One-way ANOVA with Tukey multiple comparison test (female) or Kruskal-Wallis with Dunn’s multiple comparison test (male). * p < 0.05 ** p < 0.01.

**ED Fig. 13.**
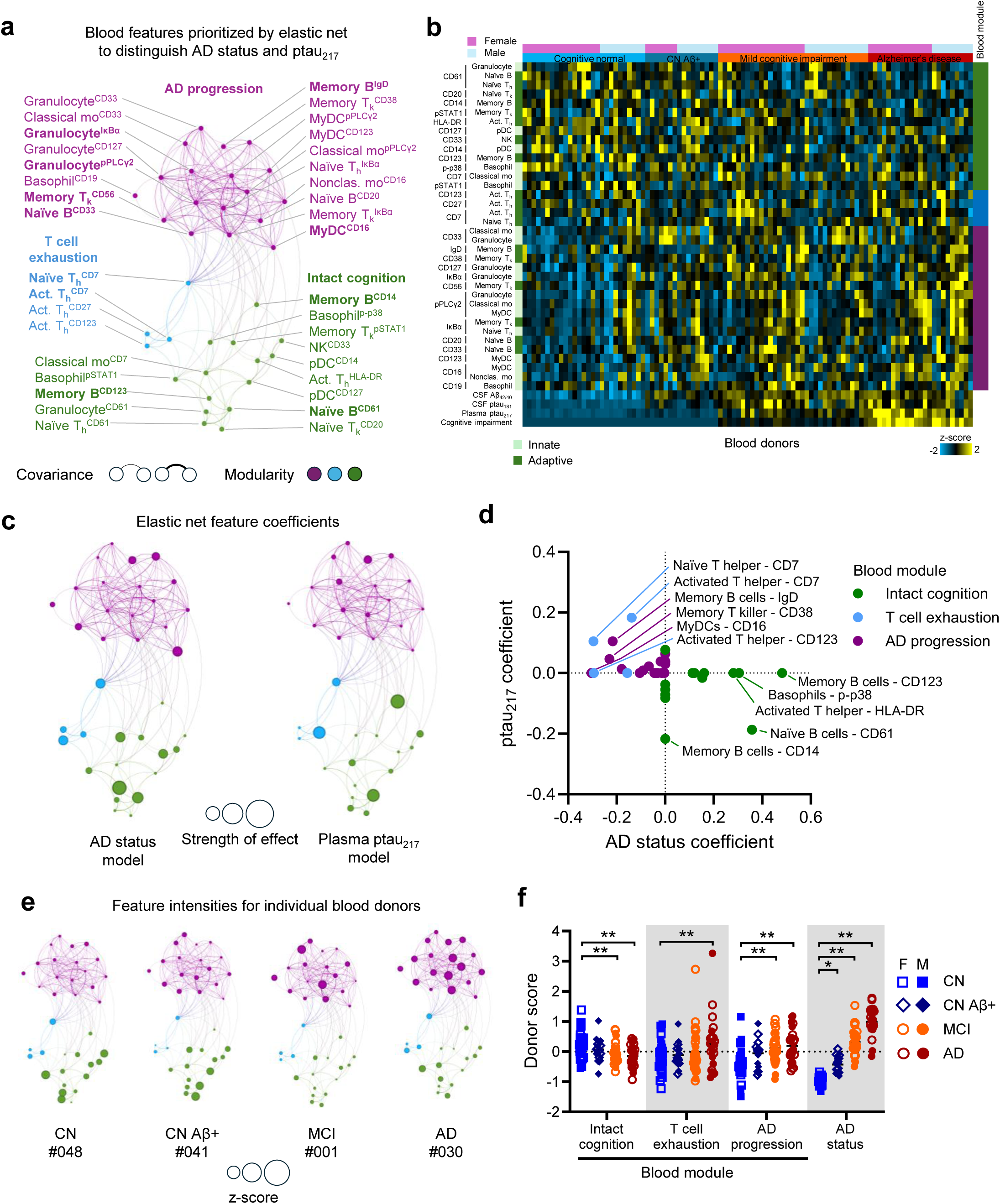
Robust detection of peripheral immune features related to ptau_217_ and Alzheimer’s disease clinical status using elastic net regularized regression. **a**, Network of peripheral blood immune features prioritized by elastic net colored by module with connections representing the strength of feature covariance. **b,** Intensity of features across female and male blood donors colored by z-score normalized intensity (teal low; yellow high). **c -d**, Network node sizes were scaled by elastic net feature β coefficient intensity (**c**) and plotted in XY graph (**d**) colored by modularity. **E**, Network node sizes were scaled by feature intensity for representative individuals classified by disease stage and patient identifier number. **f**. Mean module intensity score plotted per blood donor. Each dot represents one individual. Two-way ANOVA with Dunnett’s multiple comparison test relative to CN. * p < 0.05 ** p < 0.01.

